# Role of Chloroplast Lipid-Remodelling Protein 23 During Cold Acclimation in *Arabidopsis thaliana*

**DOI:** 10.1101/2025.07.22.666119

**Authors:** Wing Tung Lo, Denise Winkler, Maximilian Münch, Martin Lehmann, Kira Steiner, Bettina Bölter, Cornelius Gamb, Cecilia Tullberg, Carl Grey, Tatjana Kleine, Eslam Abdel-Salam, Katharina W. Ebel, H. Ekkehard Neuhaus, Deren Büyüktaş, Sophie de Vries, Hans-Henning Kunz, Dario Leister, Serena Schwenkert

**Author notes:** Corresponding author: Serena Schwenkert. The author responsible for distribution of materials integral to the findings presented in this article in accordance with the policy described in the Instructions for Authors (https://academic.oup.com/plphys/pages/General-Instructions) is Serena Schwenkert.

## Abstract

Cold acclimation is a crucial physiological process that enables plants to adapt to low temperatures. A key aspect of this adaptation is lipid remodeling, which preserves membrane fluidity and integrity under cold stress. Proteins of the chloroplast envelope membranes are increasingly recognized for their role in acclimation to changing environmental conditions. While lipid synthesis occurs at the inner envelope membrane, little is known about specific proteins involved in lipid remodeling during cold acclimation. In this study, we investigate the role of Chloroplast Lipid Remodeling Protein 23 (CLRP23) as a component of the inner chloroplast envelope membrane. Subcellular fractionation combined with protease protection assays provided evidence for its orientation toward the intermembrane space. To explore its function, we analyzed the physiological performance and lipid composition in CLRP23-deficient mutant plants. Under cold stress, we observed significant impairments in photosynthesis and exaggerations in galactolipid response, suggesting CLRP23 is involved in lipid remodeling. Lipid overlay assays, supported by *in silico* docking analyses, demonstrate that CLRP23 can directly interact with chloroplast lipids, including galactolipids. Complementary transcriptomic and proteomic analyses reveal broader effects on cold-responsive pathways, supporting the view that CLRP23 contributes to the integration of membrane and metabolic responses during acclimation. These findings expand our understanding of protein-mediated processes during cold acclimation.

## Introduction

Chloroplasts, like all plastids, trace their origins to an ancient endosymbiotic event involving the engulfment of a free-living cyanobacteria-like prokaryote by an ancient host cell. As a result of this evolutionary history, chloroplasts are bound by a double-membrane consisting of the outer (OE) and inner (IE) envelope, which are separated by an intermembrane space (IMS; Dyall et al., 2004). In higher plants, chloroplasts not only serve as the cellular site of photosynthesis but also house the enzymatic machinery necessary for various interconnected pathways that are central to plant metabolism (Weber and Fischer, 2009).

As the interface that separates the chloroplast from the cytosol, the chloroplast envelope is a highly complex and dynamic membrane system (Block et al., 2007). The evolution of chloroplasts from ancient cyanobacteria involved an extensive transfer of genes to the nuclear genome, resulting in a strong reliance on nuclear genes and cytoplasmic machinery for protein synthesis (Leister and Kleine, 2008, Martin et al., 1998). Thus, the envelope plays a primary role in the import of nuclear encoded chloroplast proteins (Soll and Schleiff, 2004). To facilitate crosstalk between the chloroplast and the cytosol, the envelope also harbors specific protein channels and transporters to facilitate the flux of ions and metabolites (Weber et al., 2005). While the IE typically comprises α-helical transport proteins, the OE is known to harbor a number of β-barrel channels, possibly being less substrate specific as compared to the IE (Schwenkert et al., 2023a, Schwenkert et al., 2023b). Notably, only very few proteins have been described to locate to the IMS, among them monogalactodiacylglycerol synthase 1 (MGD1) and the translocon of the chloroplast IE 22 (TIC22; Vojta et al., 2007).

Beyond these functions, the envelope is also one of the major sites for plant lipid metabolism, with chloroplast membranes accounting for 80% of glycerolipids in leaf mesophyll cells (Hölzl and Dörmann, 2019). This includes the shuttling of lipids and lipid precursors, as well as the biosynthesis of chloroplast specific lipids (Hölzl and Dörmann, 2019). Several envelope proteins have been associated with chloroplast lipid metabolism, such as the IE fatty acid exporter FAX1 (Li et al., 2015), the trigalactosyldiacylglycerol complex spanning both membranes (Fan et al., 2015) and the monogalactosyldiacylglycerol (MGDG) and digalactosyldiacylglycerol (DGDG) synthases (Awai et al., 2001, Dörmann et al., 1999, Kelly and Dörmann, 2002, Shimojima et al., 1997), which are attached to the OE as well as the IE, with MGD1 facing the IMS.

The ability of plants to adjust to challenging environmental conditions is heavily dependent on changes in chloroplast metabolism. As a result, the import of nuclear encoded proteins as well as the shuttling of metabolites in and out of the chloroplast play a vital role in such acclimation processes (Kleine et al., 2021, Schwenkert et al., 2022, Eisa et al., 2020). Additionally, the chloroplast membranes undergo complex remodeling processes to maintain fluidity and stability under abiotic stress (Barrero-Sicilia et al., 2017, Degenkolbe et al., 2012, Li et al., 2020). Over the past years, proteins in the chloroplast envelope in particular have been increasingly recognized for their role in plant responses to cold stress (Trentmann et al., 2020, Schwenkert et al., 2023b, John et al., 2024, John et al., 2025).

Chloroplast Lipid Remodeling Protein 23 (CLRP23; At2g17695) was originally identified in chloroplast envelope membranes of *Pisum sativum* (Goetze et al., 2015). Due to the absence of a predicted transit peptide the protein was suggested to reside in the OE and named outer envelope protein (OEP23 – from here on named CLRP23 according to its proposed function rather than localization). Here, we show that CLRP23 localizes to the inner chloroplast envelope, binds chloroplast galactolipids *in vitro*, and is essential for maintaining photosynthetic performance and balanced lipid remodeling during cold acclimation in Arabidopsis.

## Results

### CLRP23 shares structural similarities to the SRPBCC protein superfamily and is strongly attached to the membrane

To gain insight into the structure of CLRP23 (At2g17695) and to identify conserved features, an *in silico* search using the template-based modelling software Phyre2.2 was performed (Powell et al., 2025). Intriguingly, high similarity to proteins of the SRPBCC (START/RHO_alpha_C/PITP/Bet_v1/CoxG/CalC) superfamily was found. The top three ranking protein templates for the structural prediction analysis are members of the STAR-related lipid transfer (START) domain protein family, which are known for their ability to bind hydrophobic ligands (Supplementary Fig. S1B; Radauer et al., 2008; Soccio & Breslow, 2003). Overall, 139 residues (with a coverage of 68%) were predicted with a confidence of 86.4% (Supplementary Fig. S1A). Structural prediction of CLRP23 in the AlphaFold Protein Structure Database (AlphaFold DB; Jumper et al., 2021; Varadi et al., 2023) is highly similar to the model predicted by Phyre2.2 (Fig. 1A). Unlike the classical β-barrel protein channels and α-helical transport proteins typically found in the chloroplast envelopes, CLRP23 appears to have a mixture of both α-helixes and β-sheets in its predicted structure, with the β-sheets forming an incomplete β-barrel.

**Figure 1:**
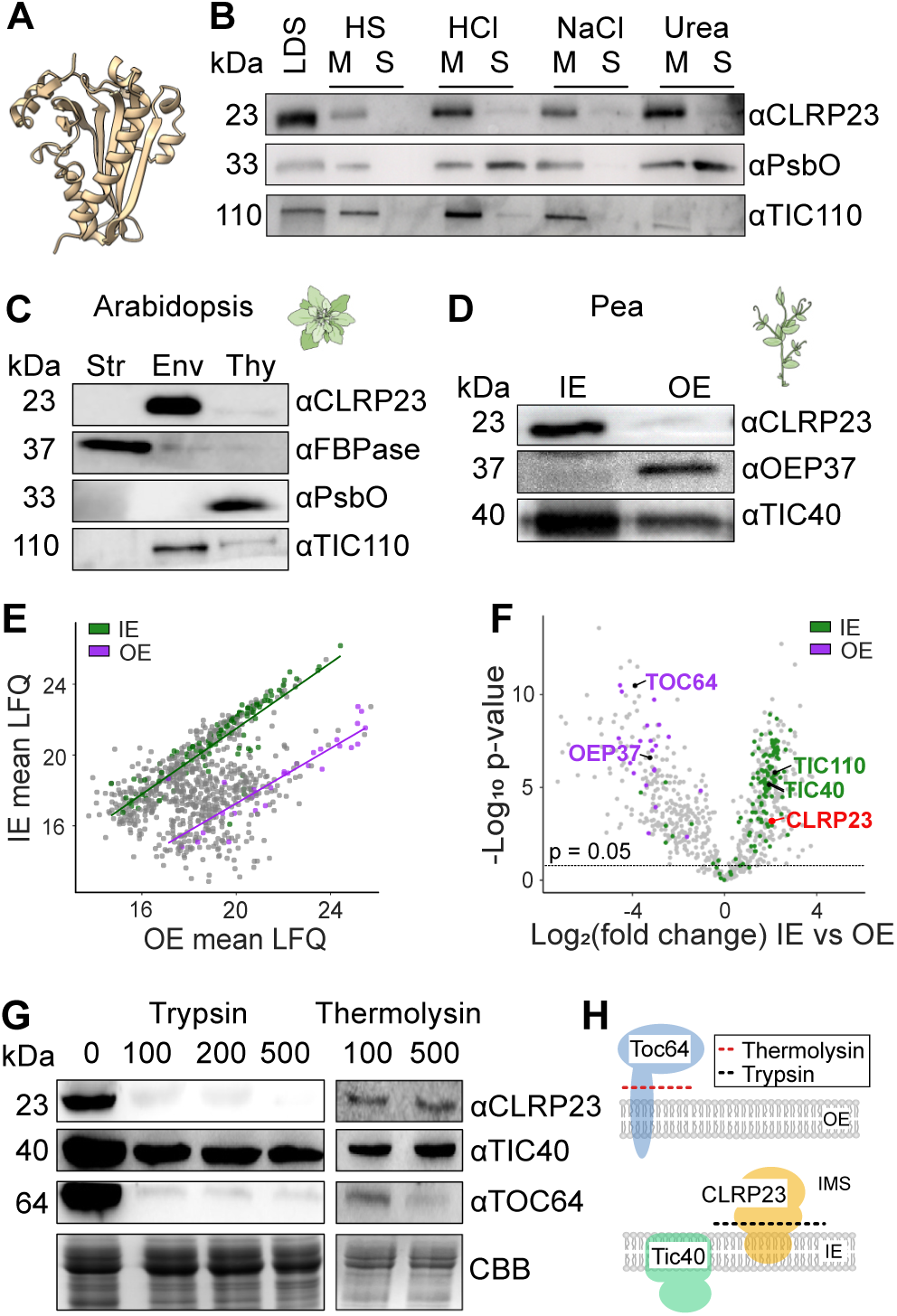
Localization and topology of CLRP23. **A)** CLRP23 AlphaFold DB structure prediction. **B)** Membrane association of CLRP23. Arabidopsis leaf tissue was treated with LDS, buffer (HS), HCl, NaCl or Urea and subsequently separated into soluble (S) and membrane (M) fractions. Samples were separated on SDS gels, transferred onto PVDF and immunolabeled with αCLRP23, αPsbO, and αTic110. **C)** Arabidopsis chloroplasts were fractionated into stroma (Str), envelope (Env) and thylakoids (Thy). Samples were separated on SDS gels, transferred onto PVDF membranes and immunolabeled with αCLRP23, αFBPase, αPsbO, αTIC110. **D)** Pea chloroplasts were fractionated into inner envelope (IE) and outer envelope (OE) transferred onto PVDF membranes and immunolabeled with αCLRP23, αOEP37, and αTIC40. **E)** Scatter plot comparing the mean LFQ intensities of proteins identified in IE and OE fractions of pea chloroplasts measured by mass spectrometry. Each point represents an individual protein, with IE proteins shown in green and OE proteins in purple. Trend lines were fitted separately for the two fractions. **F)** Volcano plot showing differentially abundant proteins in the chloroplast IE vs OE fractions based on mass spectrometric analysis. The log_2_ fold change is plotted against the -log_10_ of the p-value. Differentially abundant proteins were identified by a Student’s *t*-test (permutation-based FDR = 0.05). Proteins known to reside in the IE and OE are highlighted in green and purple, respectively. CLRP23 is labeled in red. **G)** Protease treatment of Arabidopsis chloroplasts. Intact chloroplasts of WT were treated with 0, 100, 200, 500 or 1000 µg Trypsin or 100 or 500 µg Thermolysin/mg chlorophyll. Samples corresponding to 15 µg chlorophyll were separated on SDS gels, transferred onto PVDF and immunolabeled with αCLRP23, αTic40, and αToc64, respectively. Molecular sizes are indicated on the left and Coomassie brilliant blue staining (CBB) is shown as a loading control. **H)** Schematic showing CLRP23 topology as determined in (G). The protease treatment and expected cleavages of Toc64 and CLRP23 is shown as dotted lines. Topology of CLRP23 facing the intermembrane space is depicted (IMS).

Next, we aimed to characterize the association of CLRP23 to the membrane. Isolated Arabidopsis membranes were treated with acidic or chaotropic reagents, total membrane and soluble fractions were obtained by centrifugation and analyzed by SDS-PAGE and immunoblotting. CLRP23 did not dissociate from the membrane fraction in the presence of HCl, NaCl or urea, indicating a strong attachment to the membrane (Fig. 1B). Antisera against photosystem II extrinsic protein O (PsbO) and the translocon of the chloroplast IE 110 (TIC110) were used as markers for peripheral and integral membrane associations, respectively. Treatment with Lithium dodecyl sulfate (LDS) was used as a positive control and released all three proteins from the membrane as expected.

### CLRP23 localizes to the chloroplast inner envelope facing the intermembrane space

Although we could confirm a strong membrane association of CLRP23 with the membrane fraction, its localization to the OE has yet to be experimentally confirmed. First, a CLRP23-GFP fusion protein was transiently expressed in tobacco leaves and imaged using laser scanning confocal microscopy (Supplementary Fig. S1C). Although some protein aggregation occurred, a clear pattern encircling the chloroplast was evident, resembling that of the OEP37-GFP control, a previously described chloroplast envelope marker (Goetze et al. 2006).

To further investigate the localization and topology of the protein biochemically in Arabidopsis as well as in pea, antisera against both proteins were generated. For subfractionation analysis, both Arabidopsis and pea chloroplasts were isolated and fractionated into stroma, thylakoid and envelopes. A sufficient separation of OE and IE membranes is only achievable in pea (Bölter et al., 2020), whereas in Arabidopsis only a mixed envelope fraction can be isolated. Accordingly, immunoblots were performed of isolated chloroplast stroma (Str), envelope (Env) and thylakoid (Thy) subfractions in Arabidopsis (Fig. 1C), as well as isolated IE and OE subfractions in pea (Fig. 1D). While we were able to confirm the localization of CLRP23 to the Env subfraction in Arabidopsis, psCLRP23 could only be detected in the IE subfraction in pea, in contrast to the prior assumption (Goetze et al., 2015). Antisera against fructose-1,6-bisphosphatase (FBPase, Str), PsbO (Thy), TIC110 (Env), OE protein 37 (OEP37, OE), and the translocon of the chloroplast IE 40 (TIC40, IE) were used as markers for each subfraction.

To assess the purity of our OE and IE pea membranes, proteins of 5 (OE) and 6 (IE) biological replicates were analyzed by mass spectrometry and quantified by label-free quantification (LFQ). In total 858 proteins were identified. To allow a better annotation of the proteins and connection to known functions and localizations all pea proteins were blasted against the Arabidopsis proteome. To identify Arabidopsis homologs, Arabidopsis gene IDs and their confirmed localizations are shown alongside the identified pea proteins in Supplementary Table S1. To assess the purity of the fractions, we compared the LFQ intensities of proteins found in the IE and OE fractions. A scatter plot of mean LFQ intensities (Fig. 1E) revealed a distinct separation between proteins annotated as IE and OE, indicating successful fractionation. Proteins associated with the IE exhibited systematically higher intensities in the IE fraction, while OE proteins were more abundant in the OE fraction, supporting the expected localization. Proteins clustering between the two groups likely represent contaminants from other chloroplast compartments. OE markers were on average enriched (log₂ fold change = 3.22) in the OE fraction relative to the IE fraction, while IE markers were enriched (log₂ fold change = 1.94) in the IE fraction relative to the OE fraction, indicating partial cross-contamination but substantial enrichment of the respective membrane proteins. To show statistically significant differential enrichment of proteins in either fraction, a Student’s *t*-test was performed and the log_2_ fold change was plotted against the –log_10_ p-value (Fig. 1F). Proteins assigned as IE or OE proteins are indicated. CLRP23 was clearly enriched in the IE fraction, with a log_2_ fold change of 2.04 when comparing IE vs. OE (Fig. 1F, Supplementary Table S1).

To further probe the localization and topology of CLRP23, a “dual protease” approach was used. Intact chloroplasts of Arabidopsis were treated with increasing amounts of Trypsin or Thermolysin, followed by immunoblotting with antisera against CLRP23 (Fig. 1G). Under the proper reaction conditions, Thermolysin digests proteins facing the cytosol while leaving the OE intact, whereas Trypsin can partially disrupt the OE and thus digest proteins that are facing the intermembrane space (IMS) (Froehlich, 2011). CLRP23 was found to be digested by Trypsin, but not Thermolysin, thus suggesting that it is localized to the chloroplast IE facing the IMS (Fig. 1H). Antisera against TIC40 and TOC64 were used as markers for IE and OE proteins, respectively. Coomassie Brilliant Blue (CBB) was used as a loading control.

### Loss of CLRP23 impairs photosynthetic performance under low-temperature stress

To elucidate the molecular function of CLRP23, several independent Arabidopsis knock-out mutants using two guide RNAs were generated using CRISPR-Cas technology and verified with Sanger sequencing to confirm homozygous gene disruption (Supplementary Fig. S2). Two lines were chosen for further analysis (Figure 2 and Supplementary Fig. S3A). To verify the knock-out on protein level immunoblotting of total protein extract with specific CLRP23 antisera was performed (Fig. 2A).

**Figure 2:**
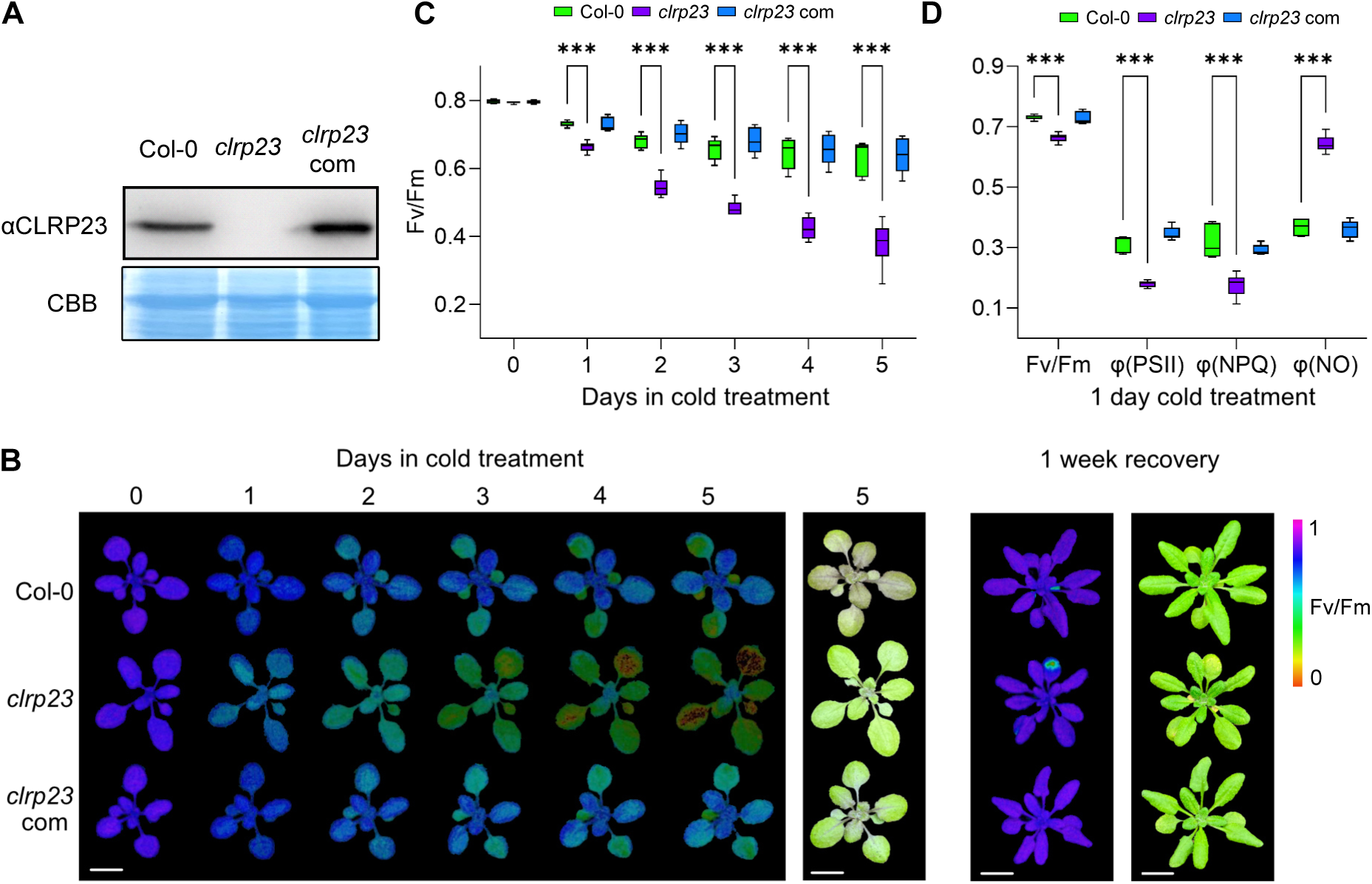
Cold sensitive phenotype of *clrp23* mutants. **A)** Total protein extracts from Col-0, *clrp23* and complemented plants (*clrp23* com) were separated with SDS-PAGE, transferred onto a PVDF membrane and immunolabeled with αCLRP23. with Coomassie brilliant blue is shown as a loading control (CBB). **B)** Imaging PAM images showing Fv/Fm and plant phenotypes. Col-0, *clrp23* and *clrp23* com were grown under standard growth conditions (SGC) for 21 days before transferring to cold treatment (4°C, 250 µE m^-1^ s^-1^). After six days of treatment, plants were transferred back to SCG for one week to recover. Scale bar = 1 cm. Chl *a* fluorescence measurements were taken from 0-5 days of cold treatment and one week after recovery. **C)** Fv/Fm values after 0-5 days of cold treatment (n=6). Statistically significant differences according to Student’s *t*-test are indicated (*p<0.05; **p<0.01; ***p<0.001). **D)** Fv/Fm, PSII quantum yield [φ(PSII)], non-photochemical quenching [φ(NPQ)] and nonregulated energy dissipation [φ(NO)] after 1 day of cold treatment (n=6). Statistically significant differences according to Student’s *t*-test are indicated (*p<0.05; **p<0.01; ***p<0.001).

In a previous proteomics study, CLRP23 was identified as differentially expressed in Arabidopsis chloroplast envelopes after cold treatments (Trentmann et al., 2020). This prompted us to investigate if *clrp23* mutants exhibit phenotypic alterations when subjected to low-temperature stress conditions. 20-day old plants grown under standard growth conditions (SGC, 21°C, 120 µE m^-2^ s^-1^) were subjected to six days cold treatment (4°C, 250 µE m^-2^ s^-1^). While Col-0 typically starts to accumulate anthocyanins after several days of cold treatment, *clrp23* did not show this behavior (Fig. 2B). To analyze the photosynthetic performance Chl *a* fluorescence parameters were measured, and *clrp23* showed a significant loss in PSII efficiency (Fv/Fm) after just one day of cold treatment (Fig. 2C). *Clrp23* also exhibited decreased PSII quantum yield [φ(PSII)], which was accompanied by decreased nonphotochemical quenching [φ(NPQ)] and increased nonregulated energy dissipation [φ(NO)], indicative of photosynthetic dysregulation and increased oxidative stress (Fig. 2D). After four days of cold treatment, Fv/Fm was drastically impaired in *clrp23* as well as the in the second independent mutant line *clrp23-2* (Supplementary Fig. S3A). Notably, photosynthetic parameters return to wild type-levels after recovery under SGC for five days. Cold-sensitivity in *clrp23* was not accompanied by noticeable effects on survival when subjected to freezing conditions (Supplemental Fig. S2B-C).

At low temperature, enzymatic activities in the chloroplast are limited while the light harvesting system remains unaffected. This imbalance can result in the generation of reactive oxygen species (ROS), thereby increasing the susceptibility of higher plants to light-induced damage (Miura and Furumoto, 2013). To separate the effects of light and cold, 20-day old plants were also treated with high light (21°C, 500 µE m^-2^ s^-1^) and cold/low light (4°C, 80 µE m^-2^ s^-1^) (Supplementary Fig. S3D-G). *Clrp23* mutants showed a significant loss in Fv/Fm after both treatments. While Fv/Fm recovered to wild-type like levels after three days of high light treatment, Fv/Fm continued to decrease under cold/low light treatment, albeit to a lesser extent. This suggests that the loss of photosynthetic performance is primarily driven by low-temperature stress and exacerbated by increased light intensity.

To ensure the cold-sensitive phenotype observed was indeed caused by the disruption of *CLRP23*, complementation lines were generated by expressing *CLRP23* cDNA under the control of the 35S promoter in both knockout mutants (Fig. 2A; Supplementary Fig. S3A). The phenotype of the complemented mutants (*clrp23 com* and *clrp23-2 com*) showed a complete recovery of photosynthetic parameters to wild-type-like levels under cold treatment (Fig. 2B-D).

### Transcriptomic and proteomic responses of clrp23 to low-temperature stress

To gain insight into the cellular pathways affected by the loss of CLRP23, we conducted transcriptomic (Supplementary Table S2) and proteomic (Supplementary Table S3) analyses comparing plants grown at SGC and plants after four days of cold treatment (4°C, 250 µE m^-2^ s^-1^). Differentially regulated genes (DEGs) and proteins (DEPs) after cold treatment vs SGC were identified by performing a Student’s *t*-test for both Col-0 and *clrp23*, respectively. The results of the Student’s *t*-tests are depicted in volcano plots (Fig. 3).

**Figure 3:**
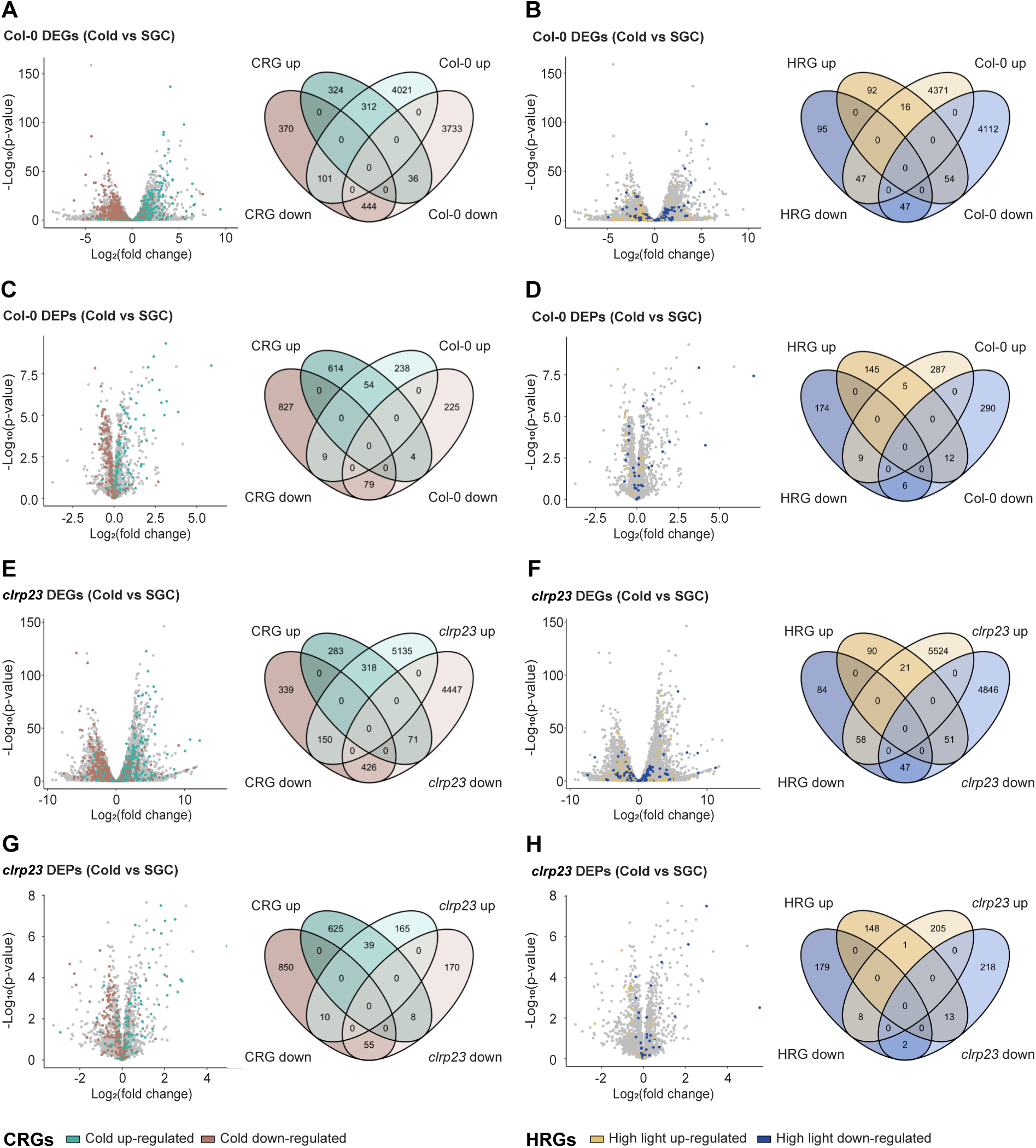
Differentially expressed transcripts and proteins in *clrp23* vs Col-0 after cold treatment. A-B) Differentially expressed genes (DEGs) in Col-0 after cold treatment (4°C, 250 µE m^-2^ s^-1^) vs standard growth conditions (SGC), in the context of published meta-analyses. DEGs were identified with DESeq2. Each dot represents one gene, plotted according to –log_10_ p-value and adjusted log_2_ fold change (n=3). Genes found in published literature to be up-or downregulated by >48 hrs cold treatment (CRGs; Hannah et al., 2005) are highlighted in green and brown (A) while genes found in published literature to be up- or downregulated by 48-120 hrs high light treatment (HRGs; Bobrovskikh et al., 2022) are highlighted in yellow and blue (B). **C-D)** Differentially expressed proteins (DEPs) in Col-0 after cold treatment (4°C, 250 µE m^-2^ s^-1^) vs SGC. DEPs were identified by a Student’s *t*-test and differentially regulated proteins were colored accordingly as above (n=4). Coloring as in (A,B). **E-F)** DEGs in *clrp23* after cold treatment vs SGC, in the context of published meta-analyses depicted as in (A,B). **G-H)** DEPs in *clrp23* after cold treatment vs SGC, in the context of published meta-analyses depicted as in (A,B).

To assess the effect of cold treatment on Col-0 and *clrp23*, we used a list of cold regulated genes (CRGs) identified in a comprehensive meta-analysis of multiple studies (Hannah et al., 2005). These CRGs were found in the meta-analysis to be differentially expressed after at least 48 h of cold exposure. CRGs detected in the transcriptomic and proteomic datasets are highlighted in the volcano plots (green for upregulated genes and brown for downregulated genes), and the overlap between CRGs and the DEGs and DEPs identified in our analyses is depicted in Venn diagrams (Fig. 3A, C, E, G). Of the overlapping CRGs, 84% in the transcriptome and 91% in the proteome of Col-0 exhibited a similar response to cold treatment compared to SGC (Fig. 3A, C). Similarly, 77% of CRGs in the transcriptome and 84% in the proteome of *clrp23* exhibited a similar response to cold as compared to SGC (Fig. 3E, G). Although *clrp23* is cold-sensitive, its transcriptomic and proteomic responses to cold are consistent with published studies on transcriptomic response to long-term cold exposure in Arabidopsis. We therefore assume it is unlikely that this cold-sensitive phenotype stems from a disruption in cold stress signaling.

A similar approach was applied to a list of high light regulated genes (HRGs) identified in a meta-analysis of high light exposure (48-120 h; Bobrovskikh et al., 2022). HRGs detected in the transcriptomic and proteomic datasets are highlighted in a separate set of the same volcano plots (yellow for upregulated genes and blue for downregulated genes), and the overlap between HRGs and the identified DEGs and DEPs is shown in Venn diagrams (Fig. 3B, D, F, H). Among the HRGs overlapping with our analysis, only 38% and 34% of the transcriptome and proteome of Col-0 exhibited a similar response under cold treatment (Fig. 3B, D). In *clrp23*, the percentages were 38% and 14%, respectively (Fig. 3F, H). These findings confirm that transcriptomic and proteomic responses to cold treatment in both Col-0 and *clrp23* appear to be primarily driven by cold.

Next, we examined the DEPs in the proteome of *clrp23* vs Col-0 under both SGC and after four days of cold treatment in more detail. While no significant alterations were observed in the proteome of *clrp23* under SGC (Fig. 4A), DEPs were identified after four days of cold treatment (Fig. 4B). Gene ontology (GO) term enrichment revealed a significant downregulation of proteins involved in flavonoid biosynthesis (Fig. 4B; Supplementary Table S4), including chalcone synthase (CHS), chalcone isomerase (CHI3), and flavanone 3-Hydroxylase (F3H), as well as enzymes acting further downstream in anthocyanin biosynthesis, including Dihydroflavonol 4-Reductase (DFR), Leucoanthocyanidin Dioxygenase (LDOX) and Anthocyanin 3-O-Glucoside Xylosyltransferase (A32gXYLT). To investigate this in more detail, anthocyanins were extracted and photometrically measured from both *clrp23* and Col-0 under SGC and after cold treatment. A reduction in anthocyanin accumulation was observed in *clrp23* after cold treatment compared to Col-0 (Fig. 4C).

**Figure 4:**
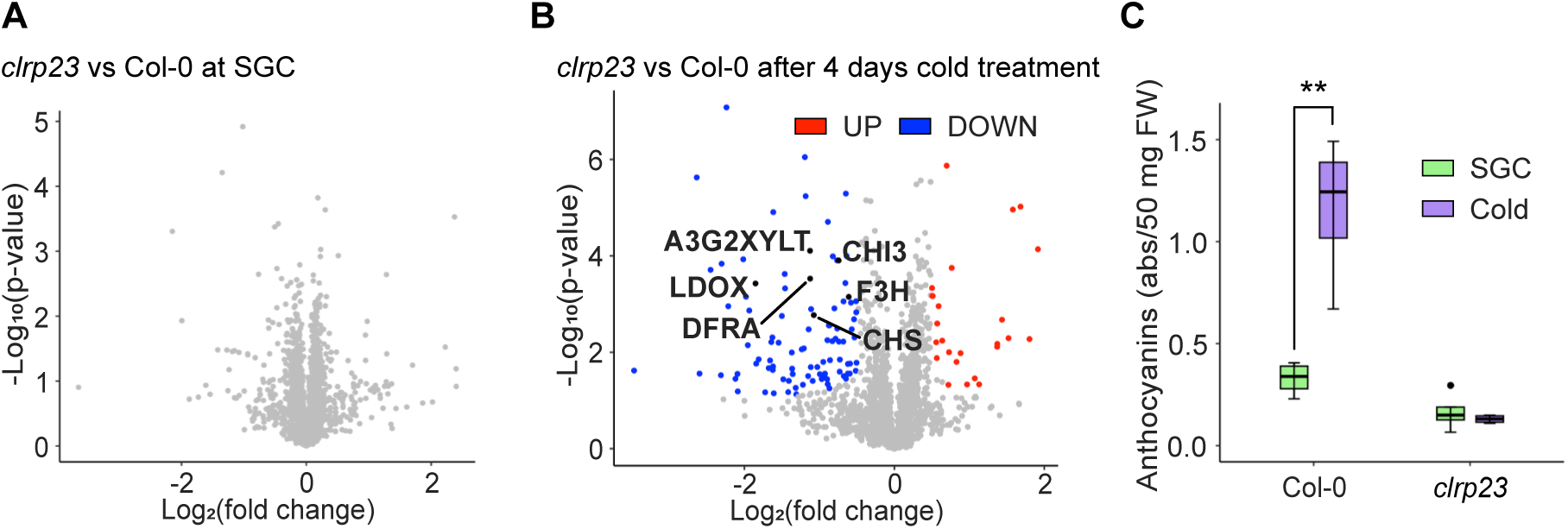
Differentially regulated proteins in *clrp23* vs Col-0. **A)** Differentially expressed proteins (DEPs) in *clrp23* vs Col-0 under standard growth conditions (SGC). Each dot represents one gene, plotted according to –log_10_ p-value and log_2_ fold change. **B)** DEPs in *clrp23* vs Col-0 after cold treatment (4°C, 250 µE m^-2^ s^-1^). DEPs were identified by a Student’s *t*-test (n=4). Significantly upregulated proteins are shown in red, downregulated proteins in blue. GO biological process enrichment analysis of DEPs in *clrp23* under cold showed a downregulation in proteins involved in flavonoid biosynthesis, the proteins of which are labeled in the volcano plot. **C)** Reduced anthocyanin accumulation in *clrp23* mutants after cold treatment. Asterisks indicate statistically significant differences according to a Student’s *t*-test (*p<0.05; **p<0.01; ***p<0.001), n=4.

### Clrp23 knockout mutants fail to accumulate neutral lipids and displays altered galactolipid remodeling under low temperature stress

Structural similarities between CLRP23 and the SRPBCC protein superfamily as shown earlier indicates the potential presence of a hydrophobic ligand binding domain. Since lipid remodeling under cold acclimation is known to play a role in maintaining membrane fluidity and influence cold signaling (Barrero-Sicilia et al., 2017, Degenkolbe et al., 2012), we investigated the effect of cold treatment on the lipid composition in *clrp23* (Supplementary Table S4). A liquid chromatography-mass spectrometry (LC-MS) based method was first used to determine the lipidomes of *clrp23* and Col-0 under SGC and after four days of cold treatment (Fig. 5A). This was complemented with gas chromatography-flame ionization detection (GC-FID) to analyze the fatty acid composition of total lipids, as well as MGDG and DGDG lipid fractions (Supplementary Fig. S4-5). Quantification of total lipids using GC-FID revealed no significant changes in total lipid content in *clrp23* and Col-0 under both SGC and after cold treatment (Fig. 5B). Z-score based k-means clustering analysis was used to group LC-MS quantified lipid data into five clusters. While most lipid classes were distributed across multiple clusters, certain lipid classes showed distinct grouping patterns (Fig. 5A).

**Figure 5:**
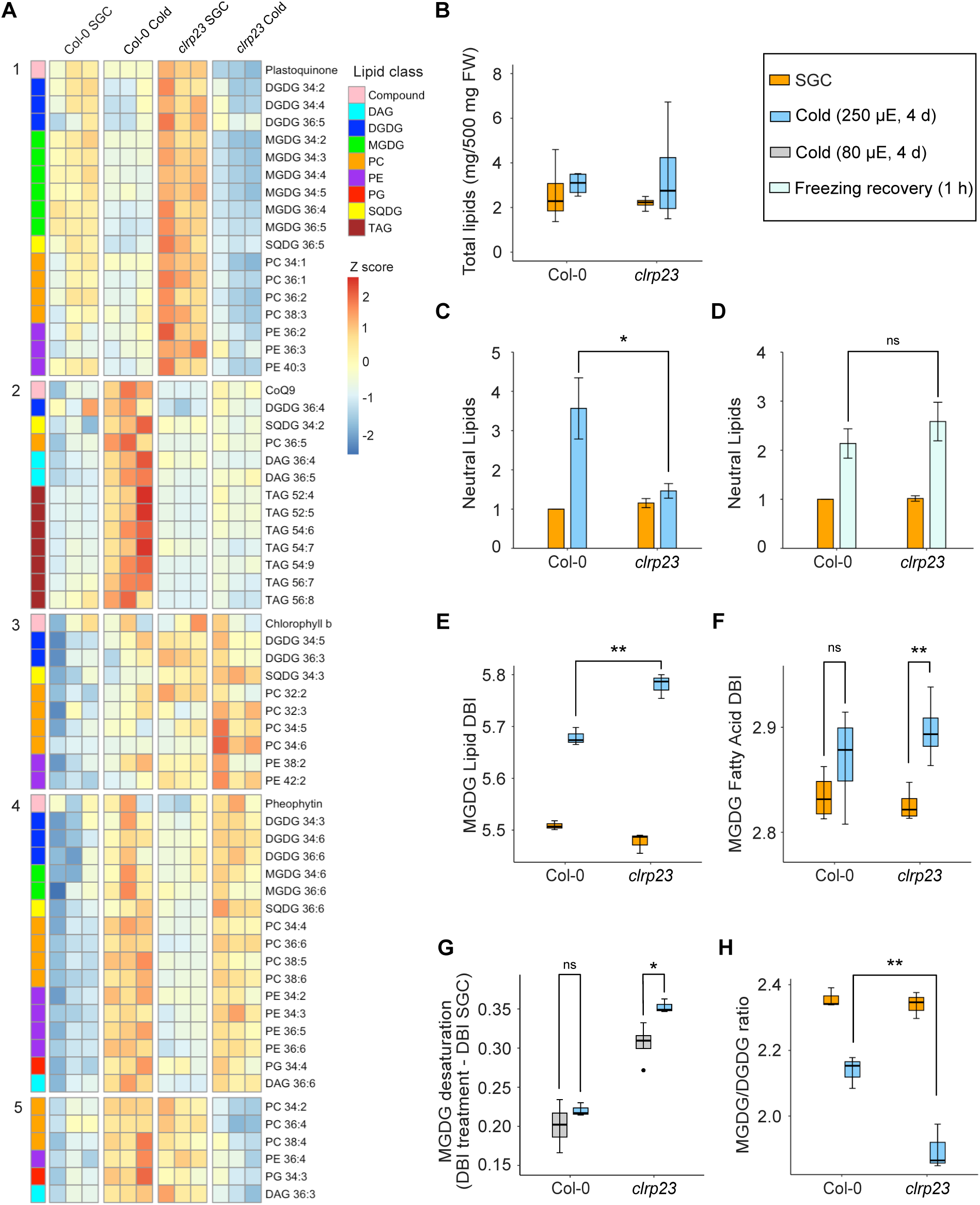
Lipid analysis of Col-0 and *clrp23* after cold treatment. **A)** Heat map showing the z scores of all lipid species detected and quantified with LC-MS in Col-0 and *clrp23* after cold treatment (4°C, 250 µE m^-2^ s^-1^) and standard growth conditions (SGC). Using k-means clustering, each lipid species is grouped into 1 of 5 clusters. **B)** Total lipids extracted and quantified with GC-FID in *clrp23* after cold treatment (n=4). **C-D)** Neutral lipids quantified with LC-MS in *clrp23* mutants after (C) cold treatment (D) and freezing recovery, reported as abundance relative to Col-0 levels at standard growth conditions. Significant differences according to a Student’s *t*-test are indicated (*p<0.05; **p<0.01; ***p<0.001), n=3 (C), n=4 (D). **E - F)** Double bond index (DBI) of MGDG (E) lipid species and (F) fatty acids in *clrp23* after cold treatment. DBI was calculated by Σ % lipid or fatty acid*the number of double bonds in each lipid or fatty acid. DBI of MGDG lipids in (E) was calculated from LC-MS data based on relative abundance while DBI of MGDG fatty acids in (F) was calculated from GC-FID data based on mol%. Significant differences according to a Student’s *t*-test are indicated (*p<0.05; **p<0.01; ***p<0.001), n=3. **G)** Change in DBI of MGDG lipid species in *clrp23* after cold treatment (4°C, 250 µE m^-2^ s^-1^) and cold low light treatment (4°C, 80 µE m^-2^ s^-1^). Significant differences according to a Student’s *t*-test are indicated (*p<0.05; **p<0.01; ***p<0.001), n=3. **H)** MGDG/DGDG ratio in *clrp23* after cold treatment. MGDG/DGDG ratio was calculated from LC-MS data based on relative abundance (peak area in relation to the internal standard corticosterone). Significant differences according to a Student’s *t*-test are indicated (*p<0.05; **p<0.01; ***p<0.001), n=3.

A large number of neutral lipids, namely two of the three diacylglycerol (DAG) species and all triacylglycerol (TAG) species were predominantly found in cluster 2 (Fig. 5A). To investigate this further, the relative abundance values of all neutral lipid species (DAGs and TAGs) quantified with LC-MS were summed and normalized to Col-0 levels under SGC. On average, cold treatment induced an approximate 3.6-fold increase in neutral lipids in Col-0, an effect that was not observed in *clrp23* (Fig. 5C). The accumulation of neutral lipids in Arabidopsis during cold acclimation has previously been reported and is speculated to result from the storage of excess fixed carbon as growth is arrested under low temperature conditions (Degenkolbe et al., 2012). However, neutral lipids can also accumulate due to adjustments in membrane fluidity and stability as part of the cold acclimation program. In particular, DAGs, which are later converted to TAGs, can be synthesized from MGDG by SENSITIVE TO FREEZING 2 (SFR2) under freezing conditions (Moellering et al., 2010). To ensure the lack of neutral lipid accumulation observed in *clrp23* under cold is not related to the SFR2 pathway, LC-MS lipid quantification was also performed on plants undergoing freezing recovery (Supplementary Table S5). Neutral lipids in *clrp23* accumulated to wild-type-like levels after 1 h of freezing recovery (Fig. 5D), suggesting that the lack of neutral lipid accumulation after cold treatment is due to the lack of fixed carbon resulting from impaired photosynthesis.

Six of the eight MGDG species grouped in cluster 1, where their levels decreased in Col-0 after cold treatment – an effect that appears to be even more pronounced in *clrp23* (Fig. 5A). The contrasting response of these MGDG species, compared to the highly unsaturated 34:6 and 36:6 species found in cluster 4, prompted us to further examine overall MGDG saturation levels. To assess this, the double bond index (DBI), calculated as Σ (% lipid or fatty acid × number of double bonds) was determined for all MGDG lipid species quantified with LC-MS (Fig. 5E), and for all MGDG fatty acids quantified with GC-FID (Fig. 5F). Cold treatment increased the DBI of MGDG lipids in Col-0, with an even greater increase in *clrp23* (Fig. 5E). A similar trend was observed in the DBI of MGDG fatty acids, where only *clrp23* showed a significant increase after cold treatment (Fig. 5F). The level of fatty acid desaturation appears to correlate with the severity of the downregulation of photosynthesis, where *clrp23* mutants exhibited a higher level of MGDG desaturation under cold (4°C, 250 μE m^−2^s^−1^) compared to cold low light (4°C, 80 μE m^−2^s^−1^), consistent with the level of photosynthetic impairment observed under each treatment (Fig. 5G).

MGDG is a non-bilayer forming lipid, with the tendency to form inverted hexagonal (HII) phases (Webb and Green, 1991). A reduction in MGDG/DGDG ratio was observed in Col-0 after cold treatment, with an even greater reduction observed in *clrp23* (Fig. 5H). This, along with the increased desaturation of MGDG lipids following cold treatment, indicates a more pronounced galactolipid remodeling response in *clrp23* under low-temperature stress.

In addition to the galactolipids, phosphatidylglycerol (PG) is a chloroplast membrane lipid which has also been linked to cold acclimation (Gao et al., 2020, Li et al., 2021). Although only two PG species were quantified in our lipidomic analysis (34:3 and 34:4), dysregulation in these two species was also observed in *clrp23* under cold (Fig. 5A).

### CLRP23 binds to chloroplast membrane lipids

Molecular docking simulations were conducted to explore potential interactions between CLRP23 and chloroplast membrane lipids. Using Autodock 4.2 (Morris et al., 2009), the 3D structure of CLRP23 was docked against the most abundant species of MGDG, DGDG, sulfoquinovosyl diacylglycerol (SQDG), PG and Phosphatidylcholine (PC) in Arabidopsis (Fig. 6). CLRP23 showed the strongest binding affinity for PG (- 6.8 kcal/mol) and MGDG (-6.7 kcal/mol), with weaker binding affinities observed for DGDG (-4.6 kcal/mol) and SQDG (-4.3 kcal/mol). To biochemically assess the potential lipid-binding function of CLRP23, we expressed it as a maltose-binding protein (MBP) fusion in *E. coli*. The soluble MBP-CLRP23 fusion protein was purified and subjected to a lipid overlay assay (Fig. 7A). MGDG, DGDG, SQDG, PG, and PC were spotted onto a PVDF membrane and incubated with the purified fusion protein. Bound protein was detected using an anti-MBP antibody. CLRP23 bound to all lipid species tested, albeit to different extents. Strong binding was observed for MGDG and PG, while interactions with DGDG and SQDG appeared weaker. Interestingly, while the strongest *in silico* binding was observed with PG, the strongest binding was observed for MGDG in the lipid overlay assay. No binding was detected for the solvent control. An MBP fusion of the unrelated chloroplast protein PALE CRESS (PAC; Meurer et al., 2017) showed no binding, confirming the specificity of CLRP23 lipid interactions and excluding artefacts from the MBP tag (Fig. 7A). Although CLRP23 also showed binding to PC in the lipid overlay assay as well as a relatively strong binding affinity *in silico* (- 6.4 kcal/mol), it is noteworthy that PC is a chloroplast membrane lipid that is only found on the outer leaflet of the OE. Given CLRP23’s localization to the IE and orientation to the IMS, it is unlikely for CLRP23 to interact with PC *in vivo*.

**Figure 6:**
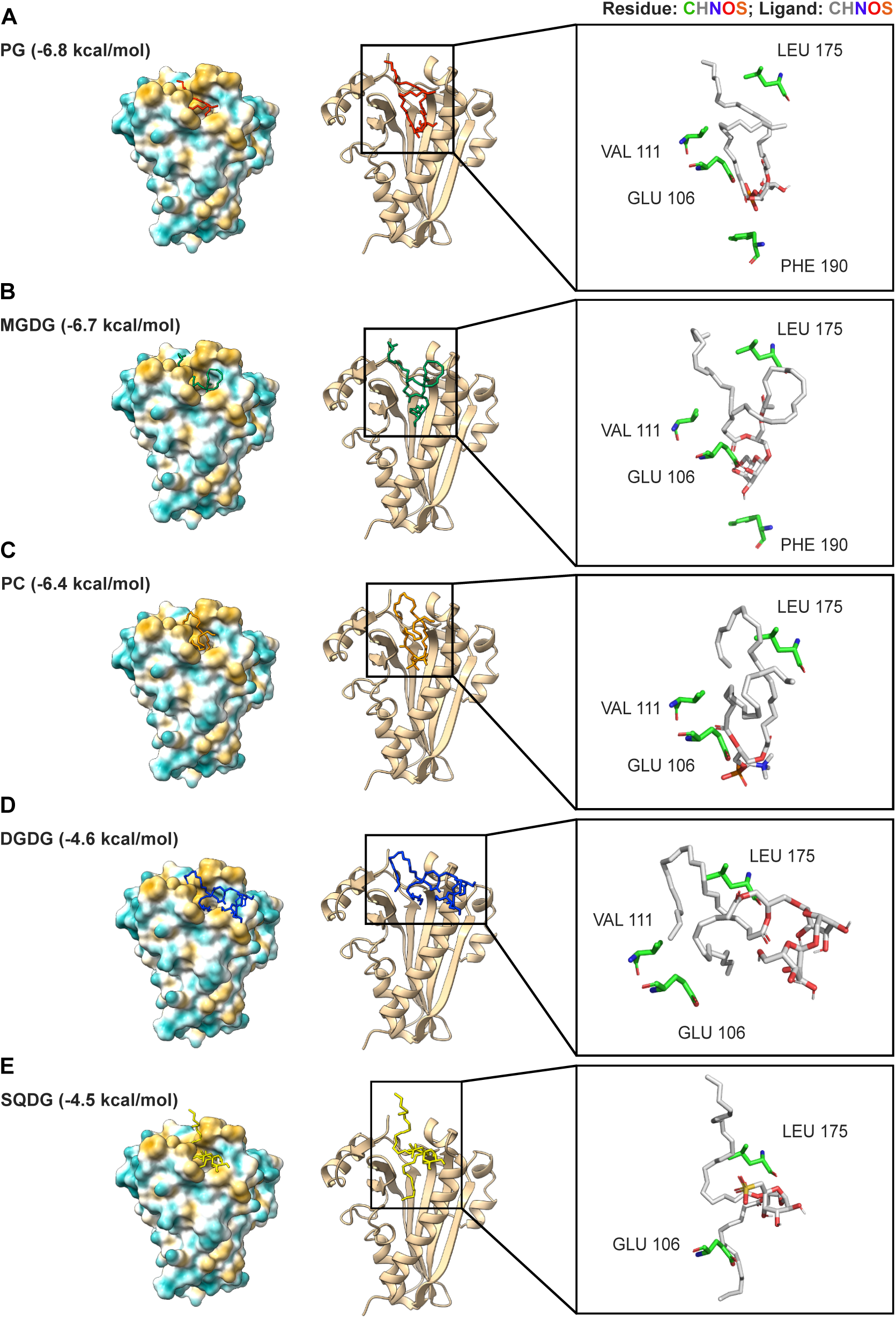
Molecular docking simulation of CLRP23 against A) PG, B) MGDG, C) PC, D) DGDG and E) SQDG. CLRP23 is depicted using a surface hydrophobicity map, colored from blue (hydrophilic) to yellow (hydrophobic), as well as a ribbon diagram. Ligand contact sites were identified with a distance-based approach (<3.5 Å) using PyMol, highlighting residues common to at least four ligands. Autodock 4.2 was used for molecular docking. ChimeraX-1.8 and PyMOL was used for visualization.

**Figure 7:**
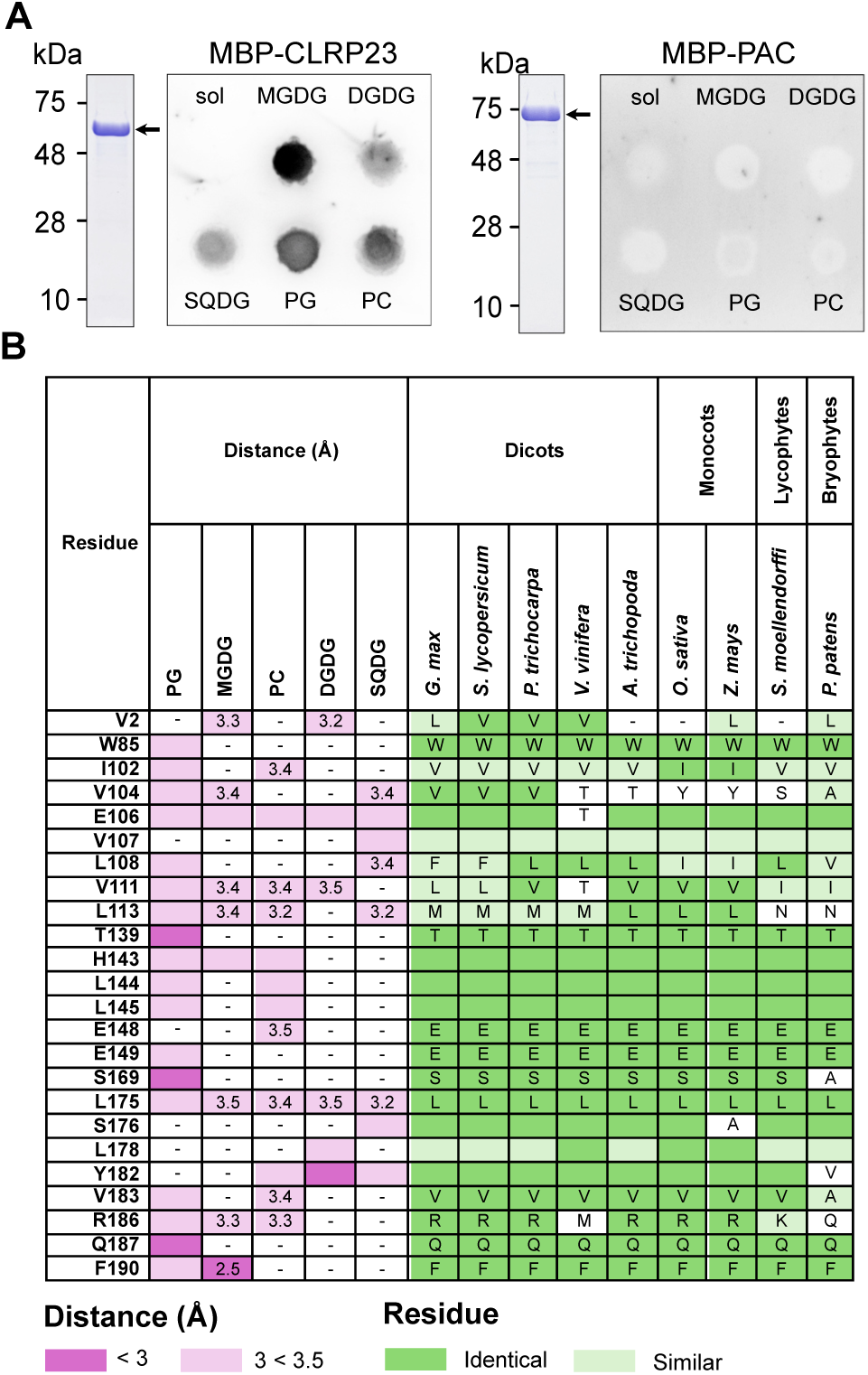
Lipid binding to CLRP23 and identification of potential binding sites. **A)** MBP-CRP23 was expressed and purified from *E. coli* (left panel). The purified protein is shown in a coomassie stained gel (CBB). A lipid overlay blot shows binding to MGDG, DGDG; SQDG, PG and PC. Sol = solvent control. MBP-PAC (right panel) was expressed, purified and subjected to a lipid overlay blot in the same manner. Arrows indicate the MPB fusion proteins. **B)** Sequence alignment analysis of homologs of CLRP23 in dicots, monocots, lycophytes and bryophytes. Homologs of CLRP23 were identified in *G. max* (Glyma.16G120900.1.p), *S. lycopersicum* (Solyc02g083180.2.1), *P. trichocarpa* (Potri.005G105500.2), *V. vinifera* (VIT_200s0338g00020.1), *A. trichopoda* (evm_27.model.AmTr_v1.0_scaffold00001.98), *O. sativa* (LOC_Os03g01730.3), *Z. mays* (Zm00001d016543_P003), *S. moellendorffi* (36490), and *P. patens* (Pp3c1_37080V3.2.p). Residues identified as contact sites for MGDG, PC, PG, DGDG and SQDG are shown as well as their respective distances to the ligand (Å). Residues in each homolog that are identical to that of At-CLRP23 are highlighted in bright green, while residues that are changed to an amino acid in the same category (hydrophobic, hydrophilic, acidic, or basic) are highlighted in a lighter green.

While all lipid ligands dock to the same hydrophobic binding pocket in the protein, the hydrophilic heads of PG, MGDG and PC are oriented inside of the pocket, while the heads of DGDG and SQDG face outwards. A distance-based approach (<3.5 Å) identified 17 amino acid residues that come into contact with at least one of the lipids in the docking simulations (Fig. 7B). To assess the conservation of these residues across species, we investigated the phylogenetic background of CLRP23. A sequence-based search for the top 100 homologs was performed against streptophytes, chlorophytes as well as the NCBI database excluding Viridiplantae (e-value <10^-7^, Supplementary Table S6). A phylogenetic analysis revealed two distinct clades: one comprising the Chloroplastida, the other consisting of prokaryotic sequences (Supplementary Figure 6). To further explore the evolutionary distribution, we searched the NCBI database for Rhodophythes, Glaucophytes and Cyanobacteria. In addition, a search excluding Viridiplantae and bacteria was performed. These searches retrieved homologous sequences from Chlorophytes, Haptophytes and Amorphaea, as well as from archaea and a limited number of cyanobacteria (Supplementary Table 6). However, homologs in Asgard archaea, the lineage that gave rise to eukaryotes (Eme et al., 2023) were not found. Next, a structure-based search was performed. With this approach, homologues were mainly retrieved from bacteria (pLDDT≥70, Supplementary Table 6, Supplementary Figure 7). Beyond the sequence-based approach, we observed a broader distribution of eukaryotic homologs, including a few Asgard archaea with a conserved core. Together, these findings suggest that the CLRP23 gene family may be ancient, although restricted to specific eukaryotic groups, a few archaea and some cyanobacteria.

Next, we aligned closely related CLRP23 orthologs from nine land plant species, spanning dicots, monocots, lycophytes, and bryophytes, to examine lipid-interacting residues in detail. (Fig. 7B). Ten residues are fully conserved among the nine homologs we analyzed. Other residues that are less well conserved often saw amino acid substitutions with similar properties. One residue of particular interest is F190, which was only identified as a contact site in the docking simulation with MGDG and PG. F190 also has a relatively short distance to the ligand (2.5 Å) and is fully conserved among the putative orthologs analyzed. These data suggest that the CLRP23 family is not only traceable to the last common ancestor of land plants but that also its lipid-interaction is conserved across land plants.

## Discussion

Prior to this study, psCLRP23/OEP23 (At2g17695) had been described as a protein channel based on electrophysiological measurements suggesting cation-selective activities *in vitro* (Goetze et al., 2015). Due to the absence of a predicted transit peptide, the protein was assumed to localize to the OE, since these proteins typically lack classical transit peptides (Kim et al., 2019). However, the localization of CLRP23 to the OE has not been experimentally demonstrated. In this study, we provide evidence that CLRP23 localizes to the IE, facing the IMS, and explore its molecular role in lipid remodeling during cold acclimation.

Based on structural predictions using AlphaFold2 and Phyre2, CLRP23 consists of a mixture of both α-helical and β-sheet structures resulting in the formation of an incomplete β-barrel. We also observed a high degree of structural similarity between CLRP23 and proteins in the SRPBCC superfamily, a protein superfamily characterized by a common 3-dimensional structure, albeit similarities on the sequence level are often less pronounced. Proteins of the superfamily typically comprise a large hydrophobic binding cavity for ligands such as membrane lipids, secondary metabolites, polycyclic aromatic hydrocarbons, and plant hormones (Radauer et al., 2008). Our subfractionation analysis very clearly showed an enrichment of CLRP23 in the chloroplast IE, rather than OE. Combined with protease treatments, our findings show that CLRP23, despite being strongly anchored to the membrane, is rather a peripheral membrane protein on the chloroplast IE, with a significant portion of the protein facing the IMS. Although the mechanism by which CLRP23 is targeted to the IMS without a predicted transit peptide remains to be elucidated in the future, the structural predictions, along with our experimental evidence, suggest an alternative role for CLRBP23, differing from its previously proposed function as an OE channel.

This alternative molecular role of CLRP23 was further explored by generating knock-out lines. Arabidopsis mutants deficient in CLRP23 suffer from impaired photosynthesis and reduced anthocyanin accumulation after several days of cold treatment. The cold-sensitive phenotype observed in *clrp23* adds to the growing body of evidence linking chloroplast envelope membrane proteins to plant acclimation (Trentmann et al., 2020, Schwenkert et al., 2023b, John et al., 2024, John et al., 2025).

Anthocyanins are a major class of flavonoids, a group of secondary metabolites characterized by their polyphenolic three-ring chemical structure (Petrussa et al., 2013). Flavonoids are primarily synthesized from photosynthetically derived carbon along the phenylpropanoid pathway and are well known to be involved in a wide variety of plant abiotic stress responses (Shomali et al., 2022, Di Ferdinando et al., 2012, Kitashova et al., 2023, Kitashova et al., 2024, Petrussa et al., 2013). During cold acclimation in particular, flavonoids accumulate after several days and have been shown to play a role in balancing C/N metabolism and maintaining protein homeostasis (Kitashova et al., 2024). However, unlike *clrp23*, photosynthesis of mutants deficient in flavonoid biosynthesis decrease in a similar manner during cold acclimation compared to Col-0 (Kitashova et al., 2023). The downregulation of flavonoid metabolism is therefore unlikely to be the sole cause of the cold-sensitive phenotype observed in *clrp23*. Conversely, it has been shown that Arabidopsis mutants suffering from impaired photosynthesis also exhibit deficiencies in the synthesis and accumulation of flavonoids (Scherer et al., 2024, Vicente et al., 2023). Thus, it is more likely that the reduced flavonoid biosynthesis and anthocyanin accumulation observed in *clrp23* mutants under cold conditions are due to a lack of fixed carbon resulting from impaired photosynthesis. Consistent with this, triacylglycerols also fail to accumulate in *clrp23* under cold conditions, suggesting a broader disruption in carbon allocation.

Beyond flavonoid metabolism, lipid remodeling is another key aspect of cold acclimation and is crucial for preserving membrane fluidity and structural integrity. The primary chloroplast membrane lipids, galactolipids MGDG and DGDG, undergo structural and compositional modifications in response to low temperatures (Barrero-Sicilia et al., 2017, Degenkolbe et al., 2012, Li et al., 2020). To counteract the rigidifying effects of low temperature, membrane lipids undergo an increase in fatty acid desaturation, introducing double bonds that create conformational kinks in lipid hydrocarbon chains. These structural changes reduce intermolecular packing and thereby enhance membrane fluidity (Hugly and Somerville, 1992, Murakami et al., 2000, Routaboul et al., 2000). In addition, MGDG is a non-bilayer forming lipid that is conical in shape with a tendency to form hexagonal HII phases (Webb and Green, 1991). To maintain membrane integrity and stability, plants also reduce the ratio of non-bilayer to bilayer forming lipids such as MGDG/DGDG as an adaptive strategy to a variety of environmental stresses, including low temperature stress (Li et al., 2020, MacDonald et al., 2023, Moellering et al., 2010, Zheng et al., 2016). Intriguingly, *clrp23* exhibited an exaggerated MGDG fatty acid desaturation as well as a more pronounced decrease in MGDG/DGDG ratio as compared to Col-0 after cold treatment, strongly suggesting an imbalance in lipid remodeling on various levels during cold acclimation (Figure 8).

**Figure 8:**
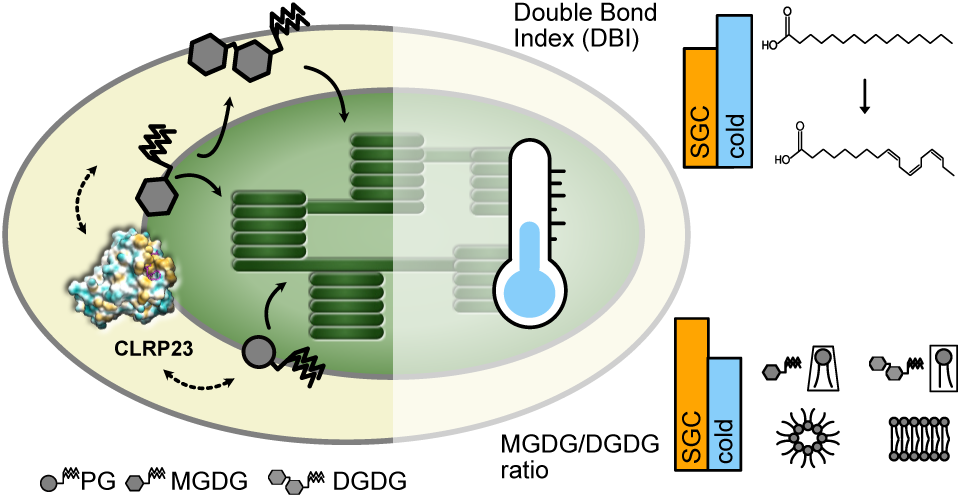
Schematic showing galactolipid synthesis at the chloroplast envelope and galactolipid remodelling under low temperature stress.

Lipid remodeling is highly complex, and involves the intricate balance between membrane fluidity, stability and functionality. While the biophysical characteristics of lipid desaturation is essential for cold acclimation, studies have demonstrated that other factors such as lipid shape also play a major role (Barkan et al., 2006, Degenkolbe et al., 2012). Saturated species of MGDG are predicted to be narrower than their more unsaturated counterparts (Gounaris et al., 1983, Gruner et al., 1985) and increases in MGDG saturation has also been postulated to help stabilize the membrane against the formation of H_II_ phases (Degenkolbe et al., 2012). A lipidomics study of 15 different Arabidopsis accessions found that the more saturated MGDG species 34:2 and 34:3 correlated positively with acclimated freezing tolerance (Degenkolbe et al., 2012). Hence, MGDG desaturation beyond what is required for membrane fluidity, such as that observed in *clrp23* under cold treatment, may have detrimental effects on membrane stability.

Furthermore, MGDG lipids make up 50% of the lipids present in the thylakoid membrane and, along with other thylakoid lipids, are integral to the assembly and function of photosynthetic complexes (Hölzl and Dörmann, 2019, Yoshihara and Kobayashi, 2022). In particular, the conical shape of MGDG has been found to sterically complement the hourglass shape of trimeric Light-harvesting complex II (LHCII), stabilizing it against unfolding (Seiwert et al., 2017). While the reduction of MGDG/DGDG ratio is a well-documented adaptive strategy against abiotic stress, Arabidopsis maintains a higher ratio than rice plants despite being more tolerant to low temperatures (Zheng et al., 2016), suggesting that the ability of plants to modulate this ratio is more crucial than its absolute value. We therefore suggest that the excessive reduction of MGDG/DGDG ratio, combined with an excessive increase in MGDG desaturation observed in *clrp23* under cold, may destabilize the membrane and alter packing relationships between membrane lipids and photosynthetic complexes, ultimately disrupting lipid-protein interactions crucial for photosynthetic function.

Changes in the lipidome, as well as the structural similarities between CLRP23 and the SRPBCC superfamily suggest a direct interaction of CLRP23 and chloroplast lipids. Indeed, lipid overlay assays supported by *in silico* docking simulations of CLRP23 revealed an affinity for chloroplast membrane lipids, with the strongest binding observed for MGDG. The residue F190, which showed the a relatively short ligand distance and was found to dock exclusively with MGDG, was completely conserved in all orthologs of CLRP23 in our sequence alignment analysis, emphasizing a potential occurrence of CLRP23-MGDG interaction *in vivo*. Strikingly, CLRP23 is one of the very few chloroplast envelope proteins currently known to face the IMS. One of the few other characterized IMS proteins is MGD1, the major chloroplast MGDG synthase (Vojta et al., 2007). This raises the question of whether CLRP23 works in concert with MGD1 to fine-tune galactolipid remodeling during cold acclimation, a question that should be addressed in future studies.

Our finding that CLRP23 likely has an ancient origin aligns with its role in lipid remodeling, a fundamental process in all organisms. It has been speculated that the eukaryotic endomembrane system originated from bacterial outer membrane vesicles (Gould et al., 2016). Interestingly, lipid binding analysis and docking simulations also revealed potential affinity for phospholipids, which may suggest a broader evolutionary function among CLRP23 homologs. The absence of sequence homologs in Asgard archaea, suggests either early acquisition via endosymbiotic gene transfer followed by losses, or lateral gene transfer among prokaryotes with subsequent transfers into eukaryotes. This is consistent with the observations that (i) the top 100 homologs to Arabidopsis CLRP23 are exclusively bacterial when searching in all datasets except Viridiplantae and (ii) bacterial homologues are predominantly recovered by structure-similarity based searches. Distinguishing between the two outlined scenarios is challenging due to the recurrent lateral gene transfers among prokaryotes. However, this could be addressed by a combined sequence-and structure-based phylogenetic approach in the future.

Together, our findings suggest that CLRP23 is involved in modulating MGDG dynamics, a crucial aspect of lipid remodeling to maintain chloroplast membrane stability and function during cold acclimation. We hypothesize that its mode of action is closely linked to the activity of MGDG synthase in the IMS, where we propose that CLRP23 may function as an MGDG-binding scaffold protein during lipid remodeling under cold acclimation.

## Material and Methods

### Plant material and growth conditions

CRISPR/Cas9 mutants of *CLRP23* were generated in Arabidopsis Col-0 background. Specific guide RNAs in the 2^nd^ exon were designed using the web server https://chopchop.cbu.uib.no/ (5’-AGTACAGAGGAGTTTCGTCT-3’; 5’-TGGTAGAGAGAGTTATGAGA-3’). Guide RNAs were cloned into lvl0 vectors resulting in ProU6 driven expression units. Using GoldenGate cloning, these lvl1 guide RNA containing vectors were assembled with a ProRPS5a driven multi-intron CAS9 (Grützner et al., 2021) and a fluorescent seed coat transformation marker (Shimada et al., 2010) into a binary plant expression vector. Arabidopsis plants were transformed using *Agrobacterium* strain GV3101 by the flower dip method (Clough and Bent, 1998). Transformed seeds were selected using the fluorescence marker. In the second-generation, knock-out mutants free of the CAS9 construct were selected by screening for non-fluorescent seeds (Ursache et al., 2021). Successful knockdown of CLRP23 was verified via immunoblotting and PCR genotyping (5’-TCACGAAGCATTGACATGTTTCA-3’, 5’-ATGGTGTTCTTGAGTTGGGG-3’). PCR genotyping was performed by amplifying *CLRP23* from extracted genomic DNA (Kotchoni and Gachomo, 2009) with Taq-polymerase PCR. PCR products were sequenced with Sanger Sequencing (LMU Faculty of Biology, Genetics Sequencing Service).

To complement the *clrp23* mutant the coding sequence of *CLRP23* including a stop codon was cloned into pB7FWG2 (Karimi et al., 2002, Plant Systems Biology, Gent) using Gateway cloning. *Clrp23* mutants were transformed with Agrobacteria strain GV3101 carrying the respective construct by the flower dip method (Clough and Bent, 1998). Transformed plants were selected by treatment with glufosinate-ammonium (BASTA). Successful knockdown of CLRP23 was verified via immunoblotting and PCR genotyping using primers specific for genomic *CLRP23* (5’-TCCTTCCCTTTCATCTTCATTTTTGG-3’, 5’-TGCAGGTTCTTCTTCTCTCTACAC-3’).

Arabidopsis plants were grown on soil in a growth chamber under long-day conditions [16 h/8 h day-night cycle, 120 μE m^−2^s^−1^, temperatures 21°C/ 18°C (light/dark)] for three weeks (SGC). For cold and light treatments plants were grown at the indicated conditions. Pea plants (*P. sativum* L., cv. ‘Arvica’, Prague, Czech Republic) were grown on vermiculite for 10 days in a climate chamber [14 h/10 h day-night cycle, 120 μE m^−2^s^−1^, temperatures 20°C/ 14°C (light/dark)]. *Nicotiana benthamiana* was grown on soil under greenhouse conditions.

Freezing treatment of Arabidopsis was performed according to Trentmann et al. (2020) with small changes. Four-week-old plants were cold acclimated for four days under short day conditions (light 10 h/4°C; dark 14 h/4°C). At the following day, the diurnal cycle was turned off and the temperature was gradually lowered from 4°C to -10°C, with a stepwise temperature decrease of 2 to 4°C per h. The freezing temperature of - 10°C was kept constant for 15 h, after which the temperature was raised again to 22°C, with a stepwise temperature increase of 2 to 4°C/h. The diurnal cycle was turned on again after one day, when the temperature was recovered back to 22°C (light 10 h/22°C; dark 14 h/18°C). Two rounds of freezing experiments were performed on a combined total of 110 Col-0 and *clrp23* mutant plants each. After freezing, samples were collected for lipidomic analysis after temperatures were recovered for 1 h at 22°C.

### Immunoblot analysis

SDS-polyacrylamide gel electrophoresis (PAGE) was performed as described by Laemmli (Laemmli, 1970). Proteins, separated via SDS-PAGE, were transferred onto PVDF membranes (Immobilon-P, Millipore, Darmstadt, Germany) via semi-dry blotting.

For CLRP23 antibody generation the coding sequence of *CLRP23* and *psCLRP23* were cloned into pET51b^+^, both proteins were purified from inclusion bodies and antisera were generated (Biogenes, Berlin, Germany).

*Agrobacterium-mediated transient expression in* Nicotiana benthamiana The coding sequence of CLRP23 and OEP37 were cloned into the binary gateway vector pB7WGF2 (Karimi et al., 2002, Plant Systems Biology, Gent). Transient expression of gene constructs was performed in leaves of 4–6-week-old *Nicotiana benthamiana* plants using *Agrobacterium tumefaciens* strain Agl1. Agrobacteria harboring the respective constructs were resuspended in infiltration medium (10 mM MgCl₂, 10 mM MES/KOH, pH 5.7, 150 µM acetosyringone) to an OD₆₀₀ of 1.0 and incubated for 2 h in the dark. Leaves were infiltrated, and protein expression was analyzed two days later. Fluorescence was observed using a Stellaris 8 confocal laser scanning microscope Imaging was carried out utilizing a Leica Stellaris 5 Confocal Laser with a supercontinuum White Light Laser and a 405 nm diode. A 63x objective with glycerol as imaging medium was used. Images were processed with the Leica LAS X software.

### Arabidopsis chloroplast isolation, subfractionation and protease treatment

Arabidopsis chloroplasts and envelopes were isolated according to Dischinger and Schwenkert (2022). For CLPR23 membrane extraction Arabidopsis leaves were homogenized in wash buffer (100 mM sucrose, 10 mM HEPES-NaOH, pH 8), filtered and centrifuged for 10 min at 10.000 *g* and 4°C. The pellet fraction was treated either with buffer (100 mM sucrose, 10 mM HEPES-NaOH, pH 8) or different chaotropic reagents (final concentration of 1 M NaCl, 0.1 M HCl, 4 M urea or 2% LDS, respectively). Samples were incubated for 30 min on ice and centrifuged for 10 min at 10.000 *g* and 4°C to separate the soluble and membrane fraction before subjection to SDS-PAGE and immunoblotting.

For protease treatment, chloroplasts equivalent to 100 µg of chlorophyll were pelleted and resuspended in 100 µl of wash buffer supplemented with 0.5 mM CaCl_2_. Trypsin or Thermolysin was added at the specified amounts per mg of chlorophyll, and the mixture was incubated at 23 °C for 40 min. To stop the reaction, 1× complete protease inhibitor (CPI) was added. Chloroplasts were then re-isolated by centrifugation and washed once with wash buffer containing 1× CPI. The final pellet was solubilized in Laemmli loading buffer supplemented with 2 M urea and 1× CPI, heated at 65 °C for 5 min, and chloroplasts equivalent to 15 µg of chlorophyll were loaded onto an SDS-PAGE gel. Immunoblotting was performed as described above.

### Chlorophyll fluorescence measurements

The photosynthetic activity of PSII was assessed by measuring chlorophyll *a* fluorescence using the HEXAGON-IMAGING-PAM system (Walz, Effeltrich, Germany). Photosynthetic parameters were calculated using IMAGING-PAM software based on the equations described by Klughammer and Schreiber (2008). Three biological replicates, two leaves each, for each genotype and condition were measured.

### Extraction and measurement of anthocyanins

Anthocyanins were extracted and measured (four biological replicates) using a method described in Kitashova et al. (2023). In short, plant material was frozen and ground in liquid nitrogen before being suspended in 1 M HCl and incubated at 25°C for 30 min on shake (800 rpm). Samples were then centrifuged at 21.000 *g* for 10 min, and the supernatant was collected before the extraction was repeated once more at 80°C for 30 min. After a second centrifugation step, the supernatant was collected and pooled for photometric quantification at 540 nm. Absorbance was normalized to fresh weight.

### RNA sequencing and data analysis

Total RNA was extracted from plants using TRIzol (Invitrogen, Carlsbad, USA) and purified with Direct-zol™ RNA MiniPrep Plus columns (Zymo Research, Irvine, USA) following the manufacturer’s instructions. RNA integrity and quality were assessed using an Agilent 2100 Bioanalyzer (Agilent, Santa Clara, USA). For RNA sequencing, ribosomal RNA was depleted to generate long non-coding RNA sequencing (lncRNA-seq) libraries, capturing both nuclear and organellar transcripts. Directional lncRNA-seq libraries were prepared, and 150 bp paired-end sequencing was performed at a depth of approximately 6 G on an Illumina NovaSeq 6000 system (Illumina, San Diego, USA) at Biomarker Technologies (BMK) GmbH (Münster, Germany) following standard Illumina protocols. Three independent biological replicates were analyzed per genotype. RNA-Seq reads were processed using the Galaxy platform (Jalili et al., 2020) as described by Tang et al. (2024), with one modification: differential gene expression of both plastid- and nuclear-encoded genes was determined using DESeq2 (Love et al., 2014) with fit type set to “parametric,” multi-mapping allowed, a linear two-fold change cutoff, and an adjusted p-value of < 0.05. One replicate of the *clrp23* mutant grown at SGC was excluded as an outlier.

### Lipidome analysis

Lipids and free fatty acids were extracted following the method described by Hummel et al. (2011). Briefly, 50 mg of each sample was extracted with 1 ml of a pre-chilled methanol:methyl tert-butyl ether (MTBE) solution (1:1, v/v) at −20°C. Internal standards were added to the extraction mixture, including 2 µg ml-1 corticosterone, 0.25 µg ml-1 ampicillin, and 1 µg ml-1 chloramphenicol. Samples were shaken for 10 min at 4°C and then ultrasonicated for an additional 10 min on ice. Subsequently, 500 µl of water:methanol (3:1) was added to the mixture, which was vortexed for 10 sec and centrifuged at maximum speed for 5 min at 4°C. Phase separation resulted in an upper organic phase containing lipids and a lower phase containing polar and semi-polar metabolites. The upper organic phase (500 µl) was collected, dried under vacuum under argon gas, and stored at −80°C until further analysis.

For LC-MS analysis, a Dionex Ultimate 3000 UHPLC system (Thermo Fisher Scientific) coupled with a timsTOF mass spectrometer (Bruker Daltonik) was employed. The dried extracts were reconstituted in 100 µl of acetonitrile:isopropanol (7:3) and 5 µl was injected onto a C8 reversed-phase column (Ultra C8, 100 × 2.1 mm; Restek) with a flow rate of 300 µl min-1 at 60°C. The mobile phases were (A) water and (B) acetonitrile:isopropanol (7:3), both containing 1% (v/v) ammonium acetate and 0.1% (v/v) acetic acid. A 26 min gradient was used, beginning at 55% B and increasing to 99% B over 15 min. This was followed by a 5 min washing step at 99% B and a return to 55% B with a 5 min equilibration phase.

For mass spectrometry, an electrospray ionization (ESI) source operated in positive mode was utilized. Nitrogen was used as the dry gas at a flow rate of 8 l min^-1^, a pressure of 8 bar, and a temperature of 200°C. Mass spectra were recorded in MS mode with a range of 50–1300 m/z, a resolution of 40,000, a scan speed of 1 Hz, and a mass accuracy of 0.3 ppm. Compounds were annotated using a targeted approach based on specific m/z values at defined retention times and their isotopic patterns. Data acquisition was performed using otofControl 6.2, with further analysis conducted using DataAnalysis 5.3 and MetaboScape 2021 software. Quantification was normalized to fresh weight and internal standards, excluding features detected in blanks or wash samples with a threshold of >10%. Three independent biological replicates were measured per genotype/condition for cold treatment, four for freezing recovery. For freezing recovery, one replicate of the *clrp23* mutant grown at SGC was excluded due to technical reasons and was replaced by taking the average value of the remaining replicates.

### Analysis of fatty acid composition

Lipids were extracted for thin-layer chromatography (TLC) and fatty acid methyl ester (FAME) analysis using a method adapted from Yu et al. (2020). 500 mg of frozen and ground plant material was ground in liquid nitrogen and suspended in 6 ml CHCl3:MeOH (1:2 v/v), with 50 µl of 5 mg ml^-1^ tripentadecanoin (TAG C15:0) added to each sample as an internal standard. Cellular debris were removed by centrifugation at 2100 *g*, 4°C for 5 min. 2 ml of CHCl3 and 1.2 ml of KCl (0.45 % v/v) was added to the resulting supernatant for phrase separation, and the lower green organic phase was collected. The solvent was evaporated by nitrogen stream and the lipids were resuspended in 250 µl CHCl3:MeOH (9:1 v/v).

For TLC, lipid samples were applied to a silica gel coated glass plate (20 x 20 cm) using an automatic TLC sampler (CAMAG® ATS 4). For each sample, a total of 8 µl of lipids were separated with acetone:toluene:water (91:30:8, v/v/v) and visualised with 0.05% primuline in acetone:water (8:2 v/v) under UV light (Hölzl and Dörmann, 2021). 4 µl each of MGDG and DGDG (2.5 mg/ml, Avanti Polar Lipids) were also applied to the plate as external standards. Water was applied to MGDG and DGDG bands to prevent particle formation, and the silica was scraped with a spatula into a fresh glass tube. The silica containing MGDG or DGDG was then suspended in 0.5 ml anhydrous methanol and sonicated for 5 min, before evaporating the methanol by nitrogen stream.

Fatty acid methyl esters (FAMEs) were generated as described in Pålsson et al. (2024), slightly modified from (Leonova et al., 2008). In short, 2 ml of MeOH:H2SO4 (98:2 v/v) was added to either 8 µl lipid extract (for total lipid analysis) or dried silica from the previous TLC step (for MGDG and DGDG analysis). 10 µl of 0.25 mg ml^-1^ heptadecanoic acid (C17:0) was also added as an internal standard for the methylation reaction. Samples were methylated in a heat block at 80°C for 45 min, after which 2 ml of water and 2 ml heptane was added for phase separation. The upper heptane phase was collected and evaporated under nitrogen gas and the FAMEs were resuspended in 50 µl cyclohexane.

The resulting FAMEs from the methylation step were then analyzed on a Trace 1300 gas-chromatography (GC) machine equipped with an ISQ single quadrupole mass spectrometer (MS), a flame ionisation detector (FID) and an auto-sampler (Thermo Scientific). FAMEs were separated on a Zebron ZB-FAME column (20m x 0.18mm i.d. x 0.15 µm film thickness). The GC oven was set to a starting temperature of 90°C, which was held for 1.5 min before ramping at 5.5°C/min to 240°C. The oven was held at 240°C for 4 min until the end of the program, with a total run time of 33 min. The MS was set with an MS line temperature and ion source temperature of 250°C, and a full scan was performed with mass range of 50-500 amu and dwell time of 0.2 seconds. FAMEs were identified based on the retention time compared with the external reference standard GLC 426 (Nu-Check-Prep) and further confirmed with MS spectra using GC-MS, after which FID was then employed for the quantification of each FAME. The FID was set to a detection temperature of 260°C. Chromeleon 7.3.1.6535 (Thermo Scientific) was used for data processing.

GLC 426 standard mix was used as the external standard to both identify the retention time and calculate the response factor for each FAME. The response factor, which accounts for the difference in detector response of each fatty acid (FA) compared to the internal standard C17:0 (IS), was calculated as follows:

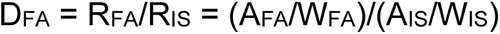

D = response factor; R = response; A = peak area (pA*min); W = weight. The response factor of C:17 was set to 1.

After subtracting the peak areas detected in the reaction blanks, the peak areas of each FAME were normalized according to the response factor of each FAME. For FAMEs C16:2 and C16:3, which were not included in the external standard, the response factor for the C16:1 FA was used for normalization. The final quantification of each FAME is reported as mol%. Total lipid content was calculated based on the sum of the normalized peak areas of all FAMEs, relative to the normalized peak area of the TAG IS C15:0, which was added as tripentadecanoin (Larodan, Sweden) to each sample prior to lipid extraction. Four independent biological replicates were used per genotype/condition for all measurements.

### Lipid overlay assay

The coding sequence of CLRP23 was amplified using oligonucleotides BamHI-OEP23-for 5’-CCAGGATCCGTGTTCTTGAGTTGGGGCCG-3’ and SalI-His-OEP23-rev 5’-CCCGTCGACTTAATGATGATGATGATGATGCGAAGCATTGACATGTTTCAGC-3’ introducing *Bam*HI and *Sal*I restriction sites as well as a C-terminal His-tag and cloned into the pMAL-TEV vector (kindly provided by Alice Barkan). Proteins were expressed in BL21 Rosetta (DE3) *E. coli* cells. MBP-CLRP23 was purified with amylose affinity chromatography (New England Biolabs) and followed by Ni-NTA-affinity chromatography (Cytivia). MPB-PAC was purified as described (Meurer et al. 2017). For the lipid overlay assay, lipids (Avanti Polar Lipids) were diluted in CHCl_3_:MeOH:H_2_O (20:10:1, v/v/v) and 7 nmol were spotted onto an activated PVDF membrane. Membranes were blocked with 0.1% (w/v) ovalbumin in TBS-T, then incubated with purified protein (1 µg/ml in 20 ml TBS-T) for 1 h at RT. Bound proteins were detected using an anti-MBP antiserum (Abcam).

### LC-MS/MS protein analysis

For Arabidopsis shotgun proteomics total proteins were extracted from 3-week-old rosette leaves by grinding frozen plant material in liquid nitrogen. A 100 mg aliquot of the resulting powder was suspended in 0.3 ml of extraction buffer (100 mM HEPES pH 7.5, 150 mM NaCl, 10 mM DTT, 6 M Guanidine Chloride, and Roche complete™ Protease Inhibitor Cocktail). The suspension was sonicated using a Branson Sonifier B-12 (Branson Ultrasonics, Danbury, USA) for three 20-second cycles and then incubated at 60°C for 10 min. Cellular debris was removed by centrifugation at 10.000 *g* for 15 min. Proteins were precipitated using the chloroform-methanol method. To the clarified supernatant, methanol, chloroform, and water were added in a 4:2:3 ratio. The mixture was centrifuged at 10.000 *g* for 10 min, and the protein interphase was collected and washed five times with methanol. The resulting protein pellet was dried and dissolved in 3 M Urea/2 M Thiourea in 50 mM HEPES (pH 7.8). Protein concentration was quantified using the Pierce 660 nm Protein Assay Kit (Thermo Fisher Scientific). For digestion, 100 µg of protein was reduced with 10 mM DTT for 30 min at 37°C, alkylated with 50 mM iodoacetamide (IAA) for 20 min at room temperature in darkness, and then digested overnight with 1 µg Trypsin (Thermo Fisher Scientific) at 37°C.

For envelope proteomics, chloroplasts were isolated from the leaves of 10-day-old pea plants and fractionated into IE and OE as described previously (Waegemann et al., 1992). IE and OE membrane samples (30 µg protein) were resuspended in 250 µl of extraction buffer, sonicated, incubated at 60°C, and centrifuged as described for plant powder. Protein digestion followed the FASP method (Wiśniewski et al., 2009). All digested peptides were purified using homemade C18 stage tips (Rappsilber et al., 2003).

For LC-MS/MS analysis, 1 µg of peptides was separated using a nano-LC system (Ultimate 3000 RSLC, Thermo Fisher Scientific) with linear gradients of 5% to 80% ACN (v/v) at a flow rate of 250 nl/min. Gradient lengths varied by sample complexity: 90 min for total proteins and 60 min for envelope samples. Chromatography was performed with Acclaim Pepmap nano-trap (C18, 100 Å, 100 µm × 2 cm) and analytical (C18, 100 Å, 75 µm × 50 cm) columns, with the column temperature maintained at 50°C. MS/MS was conducted on an Impact II high-resolution Q-TOF mass spectrometer (Bruker Daltonics, Bremen, Germany) using CaptiveSpray nano-electrospray ionization. MS1 spectra (m/z 200–2000) were acquired at 3 Hz, and the 18 most intense peaks were subjected to MS/MS at 4–16 Hz based on intensity. A dynamic exclusion duration of 0.5 min was applied.

Raw data were processed using MaxQuant (version 2.4.4.0) with peak lists matched against the Arabidopsis or *Pisum sativum* reference proteome (UniProt) using default parameters. Protein quantification was performed using the LFQ algorithm. Data were further analyzed using Perseus (version 2.0.6.0). Contaminants, reverse hits, and proteins identified only by site modification were excluded. For *clrp23* proteomics proteins quantified in at least three out of four replicates were retained. For envelope proteomics proteins quantified in at least four replicates were retained. LFQ intensities were log2-transformed, and missing values were imputed using a normal distribution in Perseus. The Database for Annotation, Visualization, and Integrated Discovery (DAVID) was used to identify enriched gene ontology (GO) terms based on biological process (Huang et al., 2009, Sherman et al., 2022).

### Molecular docking analysis

The 3D structure of At2g17695 (CLRP23) was downloaded in PDB format (UPF0548) from the AlphaFold Protein Structure Database (Jumper et al., 2021, Varadi et al., 2022). Using AutoDock 4.2 (Morris et al., 2009), the PDB files were prepared via removing water molecules, repair missing atoms, adding polar hydrogens, adding Kollman charges (Singh and Kollman, 1984), and checking total charges on residues. Several ligands were used for docking analysis, i.e., MGDG (CID: 5771744), DGDG (CID: 363608285), PC (CID: 52923341), SQDG (CID: 363648189) and PG (CID: 52926638). The 3D conformation of the different ligands was either obtained as SDF files from the PubChem database (Kim et al., 2025) or generated based on the 2D conformations SDF files using OpenBabel v3.1.1 (O’Boyle et al., 2011). SDF files were then converted into PDBQT format in the AutoDock 4.2.

All the docking simulations performed in the current study were via AutoDock 4.2 and AutoDock Vina 1.2.0 (Eberhardt et al., 2021, Trott and Olson, 2010). Default docking parameters were chosen with a globally searching exhaustiveness of 8. The docking grid size was set to 44 × 64 ×58 Å where its center corresponding to the center of the receptor protein. The docking conformation with the lowest binding affinity (kcal mol^-1^) was adopted. Molecular graphing software ChimeraX-1.8 (Pettersen et al., 2004) and PyMOL (Schrodinger, 2010) were used for further visualization and identification of contact sites.

### Sequence homology and phylogenetic analysis

The Arabidopsis CLRP23 sequence was used to query our database of land plants and streptophyte and chlorophyte algae using BLASTp (references for the sequence databases are provided in Supplementary Methods S1). The results were filtered based on e-value and hits with an e-value of ≤10^-7^ retained. Additionally, we queried the nr database with the following restriction (i) excluding Viridiplantae, (ii) excluding Viridiplantae and bacteria, (iii) Cyanobacteria, (iv) Rhodophyta and (v) Glaucophyta using an e-value cutoff of ≤10^-7^. For phylogenetic analysis the BLASTp output was aligned using MAFFT with G-INS-I (Katoh and Standley, 2013). After a quality control a maximum likelihood phylogeny was constructed using IQTree with 100 bootstraps per phylogeny (Nguyen et al., 2015). The model was selected using modelfinder implemented in the software (Kalyaanamoorthy et al., 2017). Phylogenies were visualized in iTOL (Letunic and Bork, 2021) and annotated manually for taxonomy.

Additionally, we used AlphaFold3 website (Abramson et al., 2024) for identifying structurally similar proteins in the AlphaFold3 database (AFDB50). As query we used the protein sequences of *At*CLRP23 (Q8GXB1) and investigated the taxonomic distribution across hits with a pLDDT≥70 to represent strong overall structural similarity and backbone similarity. Taxonomic identification of hits was done by recovering the TaxID for the genus a particular hit corresponded to.

To investigate residue conservation, the BLASTp search function in Phytozome 13 (Goodstein et al., 2011) was used to identify CLRP23 homologs across a list of nine land plant species spanning dicots, monocots, lycophytes and bryophytes (references for the sequence databases are provided in Supplementary Methods S1). CLC Main Workbench 20.0.4 (QIAGEN, Aarhus, Denmark) was used to conduct sequence alignment with CLRP23 in Arabidopsis.

### Statistical analysis

In addition to the software mentioned above, Microsoft Excel, GraphPad Prism, R and R studio were also used for statistical analysis and data visualization. Raw data files of transcriptomic, proteomic and lipidomic analysis can be found on DataPlant (Schwenkert et. al. 2025, doi: 10.60534/qsh62-7p088, https://www.nfdi4plants.de/).

## Acknowledgements

CRISPR/Cas9 vectors were generously provided by Marc Youles (The Sainsbury Lab, Synthetic Biology), Silvestre Marillonnet (Leibniz Institute of Plant Biochemistry Halle, Synthetic Biology), and Cyril Zipfel (University of Zurich, Molecular and Cellular Plant Physiology). We sincerely thank Benjamin Brandt for his assistance with CRISPR/Cas9 construct design. Further we are grateful to Alice Barkan for providing the MBP vector and Nikolay Manavski for providing the MBP-PAC control. We also express our gratitude to the MSBioLMU mass spectrometry service unit, particularly Yulia Davydova, for outstanding technical support. Additionally, we acknowledge Nadia Al-Thour and Tim Lücke for their valuable help with experiments. We also thank Timo Mühlhaus, David Zimmer and the DataPLANT consortium for supporting data archiving following the FAIR data principles (https://www.nfdi4plants.de/) and the Graduate School Life Science Munich (LSM) for their support. Artificial intelligence tools ChatGPT and DeepL were used to help write code and improve language. This research was funded by the German Research Foundation (DFG) TRR175, with support to SS (B06), DL (B07), HHK (B09), HEN (B03), and TK (C01). Confocal microscopy work was funded by DFG (INST 86/2231-1 FUGG) to HHK.

## Author contributions

SS designed the research and supervised the study. WTL, DW, MM, ML, TK, EA, BB, CG^1^, KS, KWE and DB performed research and analyzed data. SS, CT, CG^4^, HEN, SV, HHK and DL analyzed data. WTL and SS wrote the paper with contributions of all authors.

---

**Supplemental Figure S1:**
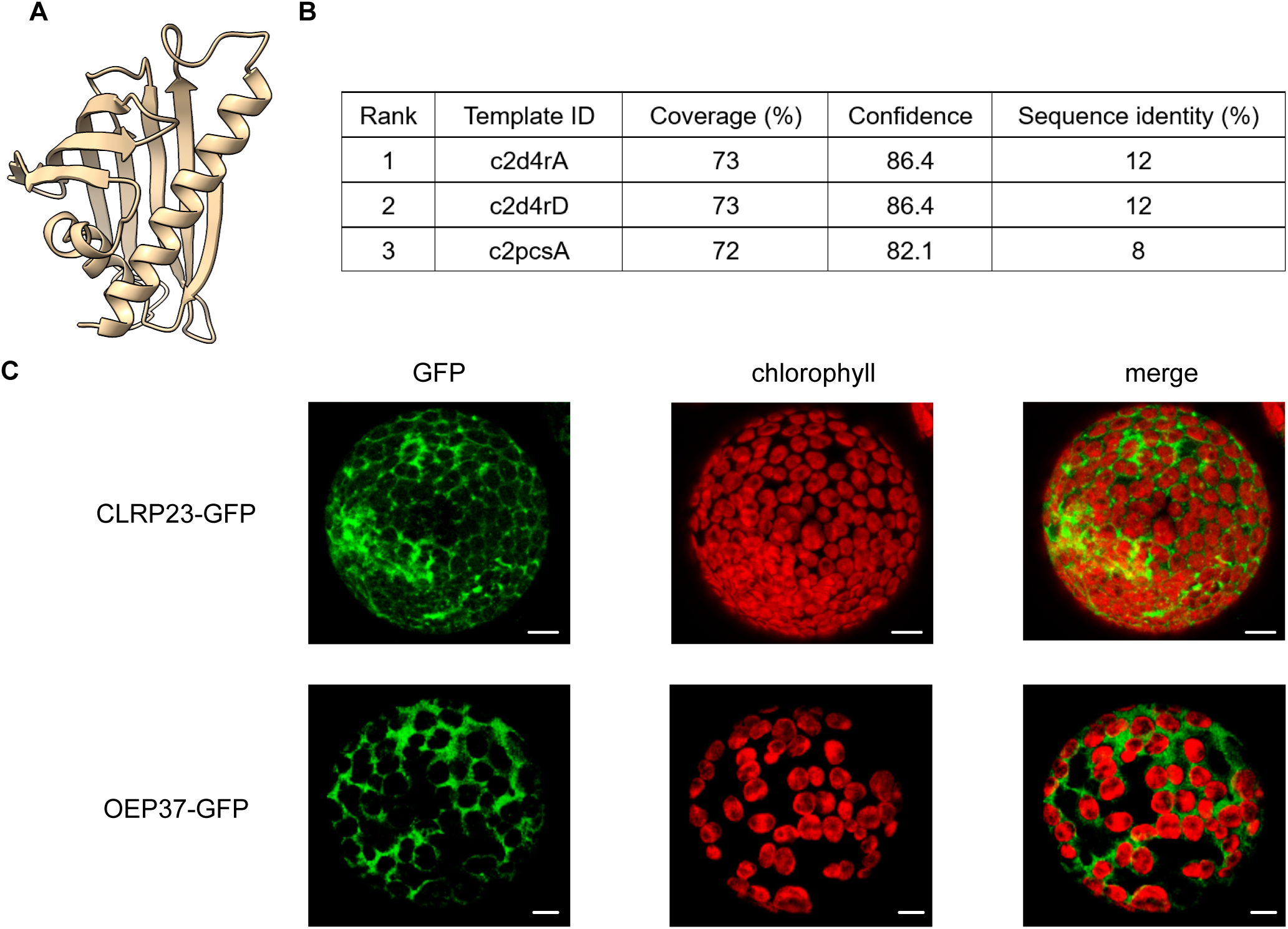
Structural prediction of CLRP23 with Phyre2.2 and GFP localisation. **A)** Visualisation of predicted structure of CLRP23 with UCSF ChimeraX-1.8. **B)** Topped ranked template IDs from structural prediction. **C)** GFP localisation of CLRP23 and OEP37. C-terminal tagged CLRP23-GFP and OEP37-GFP serving as an envelope control was transiently expressed in tobacco leaves and confocal laser scanning microscopy was performed with isolated protoplasts. A representative image showing GFP fluoresence and chlorophyll autofluorescence are shown. Scale bar =10 μm

**Supplemental Figure S2:**
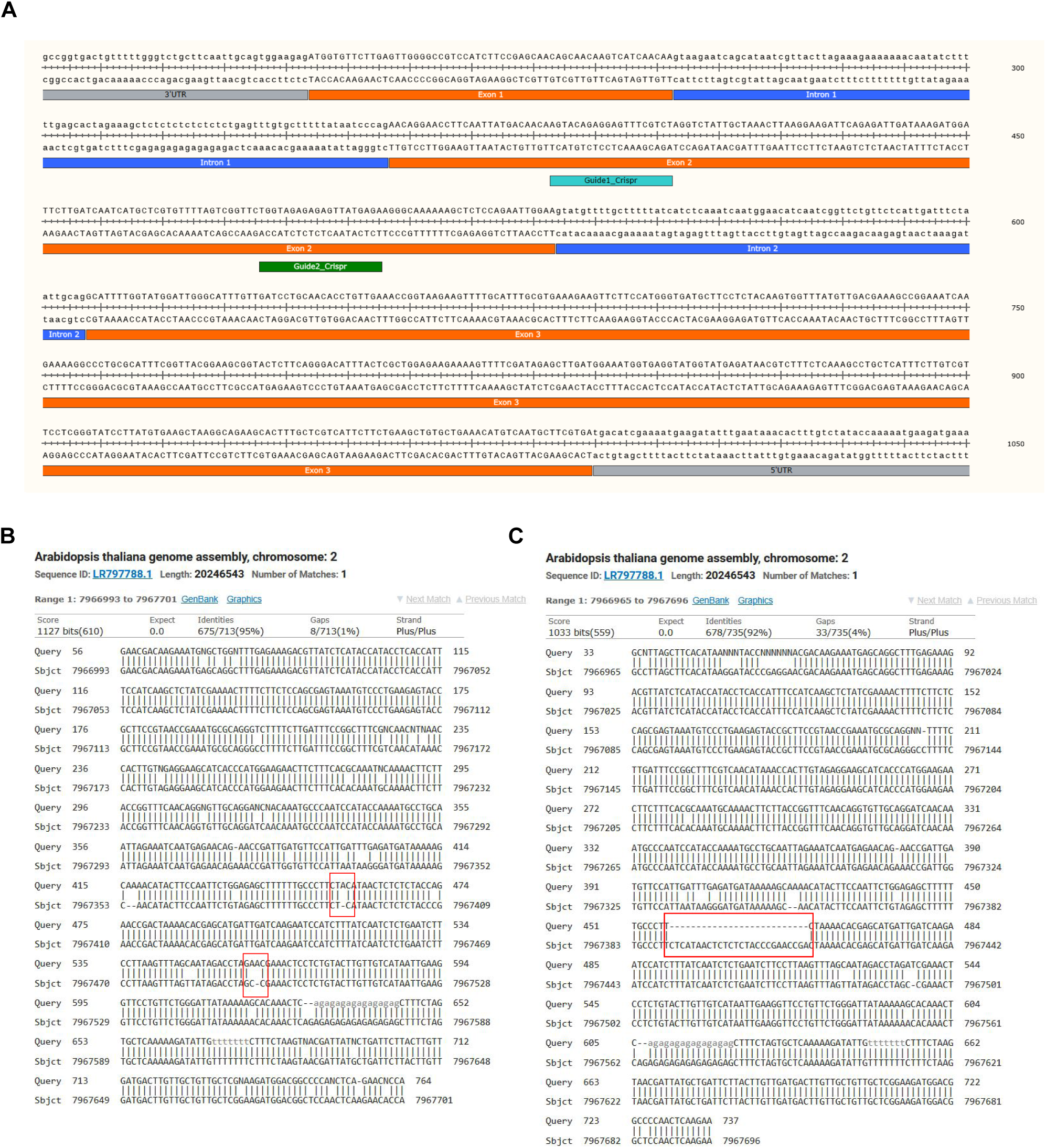
*clrp23* knock-out mutants generated with CRISPR/Cas9 technology. **A)** gDNA sequence of *CLRP23*, and the two specific guide RNAs designed for CRISPR/Cas9 mutant knock-out. **B-C)** Confirmation of gene knock-out with PCR and Sanger Sequencing of mutant lines (A) *clrp23* and (B) *clrp23-2.* NCBI nucleotide BLAST was used to align the sequencing data with the gDNA sequence of *CLRP23*.

**Supplemental Figure S3:**
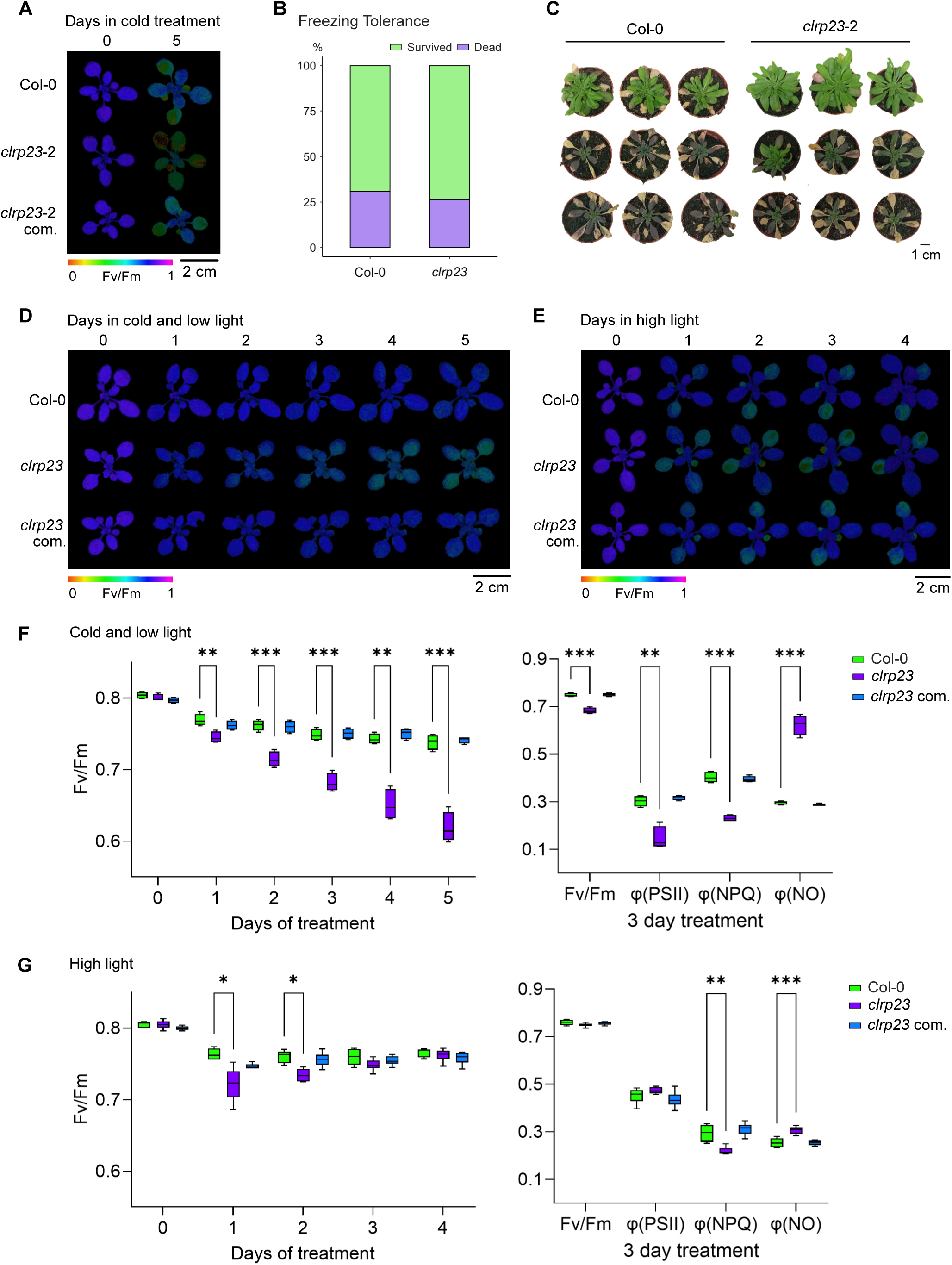
*clrp23* mutants under cold, freezing, cold, cold low light and high light conditions. **A)** Independent CRISPR mutant line lacking CLRP23, *clrp23-2* and the corresponding complementation line, *clrp23-2* com under cold treatment (4°C, 250 µE). **B-C)** Col-0 and *clrp23* plants were subjected to freezing treatment. (B) shows the survival rates of Col-0 and *clrp23* to freezing and (C) shows the representative diversity of plants responding to freezing treatment after recovery to 22°C **D-G)** Col-0, *clrp23* and complemented plants were grown under standard growth conditions for 20 days before transferring to cold/low light (4°C, 80 µE) and room temperature/ high light conditions (22°C, 500 µE). **D-E)** Chlorophyll luorescence imaging during cold low light (D) and room temperature high light (E) treatment. **F-G)** Fv/Fm values after 0-5 days, as well as Fv/Fm, PSII quantum yield (φ(PSII)), non-photochemical quenching (φ(NPQ)) and nonregulated energy dissipation (φ(NO)) after 3 days of cold low light (F) and room temperature high light (G) treatment. Asterisks indicate statistically significant differences according to Student’s t-test (*p<0.05; **p<0.01; ***p<0.001).

**Supplemental Figure S4:**
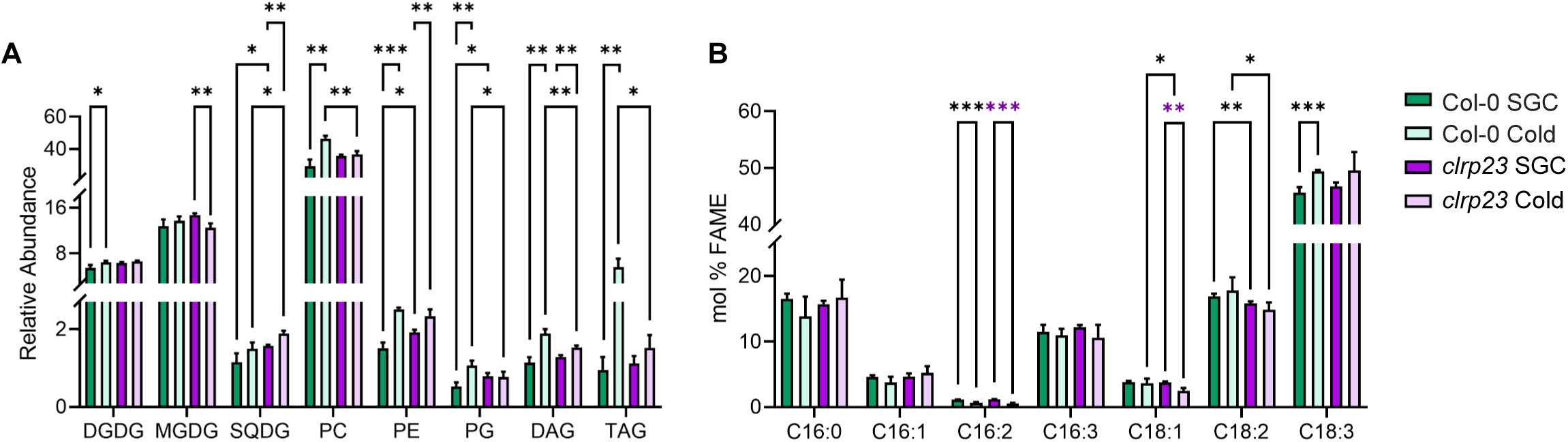
Total lipid analysis of *clrp23* mutants under standard growth conditions (SGC) and after cold treatment. **A)** Relative abundance of lipids from each lipid class summed, quantified with LC-MS. **B)** Fatty acids of total lipid extraction were quantified with GC-FID and reported as molar percentage (mol% FAME). Asterisks indicate statistically significant differences according to a Student’s t-test (*p<0.05; **p<0.01; ***p<0.001).

**Supplemental Figure S5:**
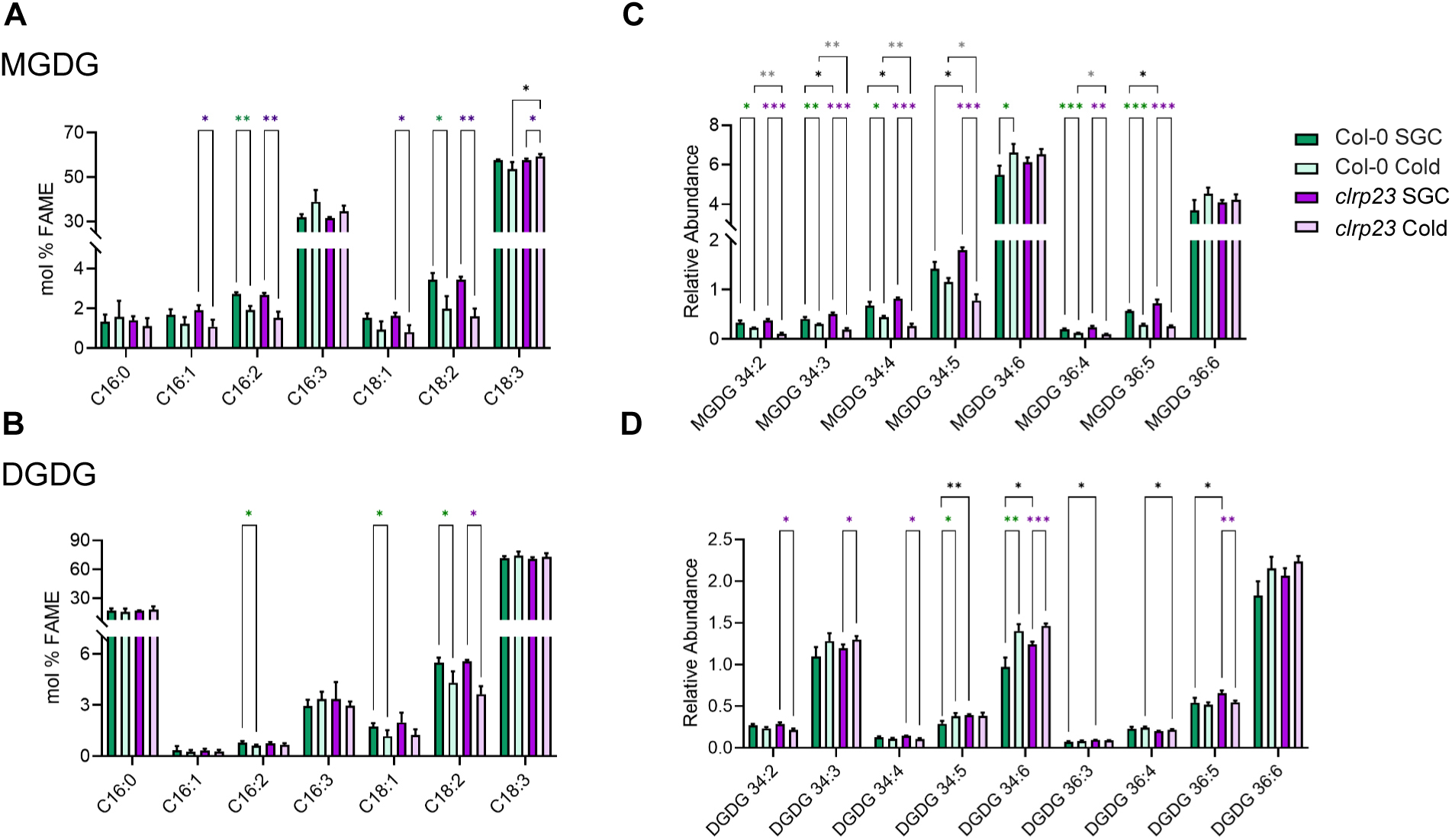
Galactolipid analysis of *clrp23* mutants after cold treatment. Side by side comparison of MGDG (A, C) and DGDG (B, D) fatty acids (A-B) and lipid species (C-D) quantified with GC-FID and LC-MS, respectively. Fatty acids are reported as molar percentage (mol% FAME) and lipid species are reported as relative abundance (peak area relative to internal standard corticosterone). Asterisks indicate statistically significant differences according to a student t-test (*P<0.05; **P<0.01; ***P<0.001).

**Supplemental Figure S6:**
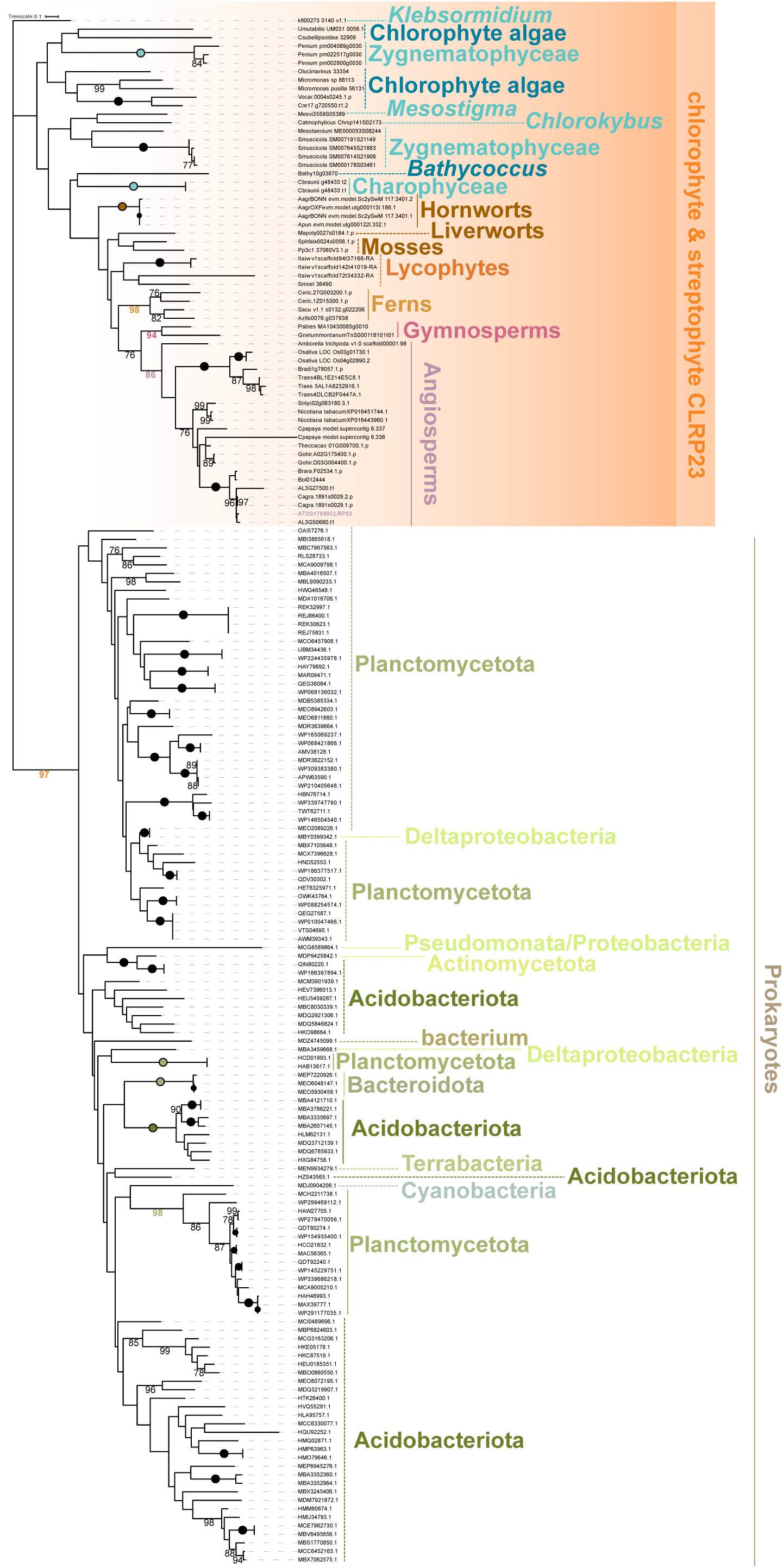
Phylogenetic analyses of CLRP23. The unrooted phylogeny is based on BLASTp hits of *At*CLRP23 against a streptophyte and chlorophyte genome containing database and the nr database on ncbi restricted to (a) bacteria and (b) cyanobacteria. *Arabidopsis* gene family member CLRP23 is highlighted in light purple and bold. The CLRP23 sequences from Chloroplastida are highlighted by an orange box. Taxonomic ailication is indicated on the right. Bold lines show monophyletic groups with a minimum bootstrap support of >75. Dotted lines indicate sequences belonging to the same lineage, which are either paraphyletic or not supported to form a monophylum. Only bootstrap support of >75 is shown, bootstrap support of 100 is indicated by circles.

**Supplemental Figure S7:**
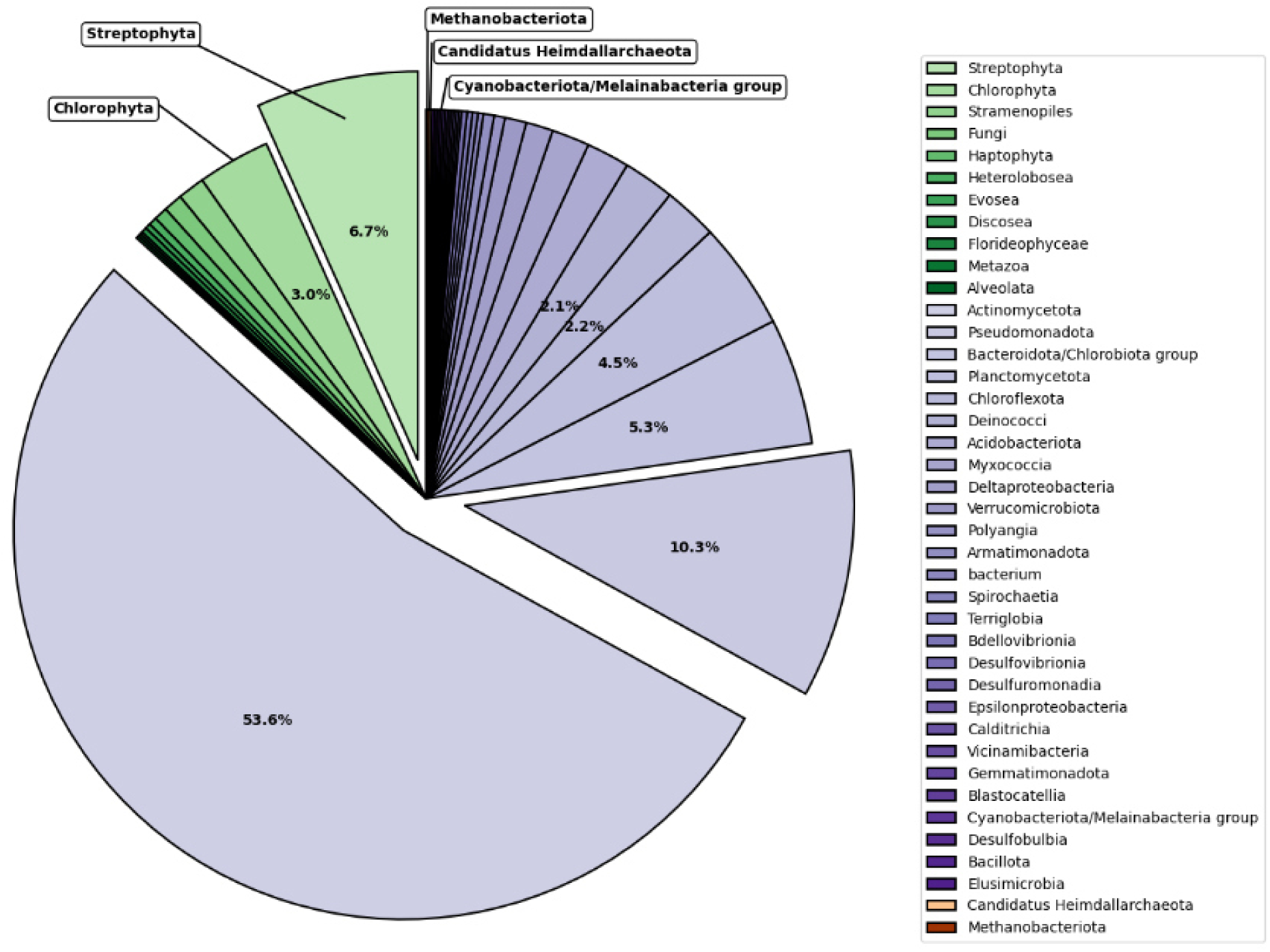
Taxonomic distribution of sequences that are structurally similar to *At*CLRP23, showing eukaryotes (green), bacteria (purple) and archaea (orange). Figure is based on results from Alphafold3 search in Table S6.

